# Cumate-inducible models of MYC-driven neuroblastoma identify mitoribosomal synthetic lethality as a precision oncology vulnerability in *MYC(N)*-amplified tumors

**DOI:** 10.64898/2026.05.25.725359

**Authors:** Adela Kubistova, Iryna Horak, Tomas Barta, Marie Sulova, Martin Marek, Jaroslav Sterba, Karolina Borankova, Jan Skoda

## Abstract

Oncogenic MYC transcription factors profoundly rewire cellular programs, imposing therapeutically exploitable dependencies in MYC-driven cancers such as high-risk neuroblastoma. However, dissecting these synthetic lethal vulnerabilities using tunable gene expression remains challenging. Widely used tetracycline-regulated systems rely on inducers known to perturb mitochondrial function, introducing confounding off-target effects. To overcome this limitation, we established novel cumate (p-isopropylbenzoate)-inducible neuroblastoma models that enable physiologically unbiased regulation of *MYC/MYCN* expression. Functional validation demonstrated that cumate alone has no off-target effects on neuroblastoma cell viability, mitochondrial function, morphology, proteostasis, or stress signaling. The developed SHEP-CuO-MYC and -MYCN models show efficient, titratable, and reversible upregulation of c-MYC and N-MYC, respectively, recapitulating protein levels observed in *MYC(N)*-amplified neuroblastoma. As a proof-of-concept, we applied these models to mechanistically validate the recently proposed mitoribosomal synthetic lethality, providing clear evidence that *MYC(N)* activation sensitizes neuroblastoma cells to mitochondrial ribosome inhibitors. While impairing mitochondrial translation activated mitochondrial integrated stress response in both MYC-on and MYC-off states, it induced marked MYC downregulation coupled with enhanced caspase-dependent cell death only in MYC-on cells. Importantly, these MYC-selective effects were reproduced in patient-derived neuroblastoma cell lines, supporting the translational potential of mitoribosomal synthetic lethality. Together, our findings reveal that *MYC(N)* activation confers a clinically relevant mitochondrial vulnerability that can be selectively targeted by repurposing doxycycline or other well-tolerated mitoribosome-targeting antibiotics. This study establishes mitoribosomal synthetic lethality as an actionable precision oncology strategy and provides a robust, scalable platform to identify MYC-synthetic lethal candidates for rapid translation into therapies for refractory MYC-driven neuroblastoma.

**HIGHLIGHTS:**

- Cumate-inducible MYC models enable unbiased studies of mitochondrial synthetic lethality
- MYC overexpression drives neuroblastoma sensitivity to mitochondrial translation inhibition
- Context-specific MYC downregulation links mitochondrial stress to MYC synthetic lethality
- Common ribosomal antibiotics trigger caspase-dependent cell death in MYC-driven tumor cells
- MYC-selective lethality is reproduced in patient-derived neuroblastoma cells

## 1 INTRODUCTION

Neuroblastoma is a malignant pediatric tumor arising from the developing sympathetic neural system [1–3]. It accounts for ∼8% of all childhood cancers [4] and nearly 15% of cancer-related deaths in children worldwide [5, 6], making it one of the most lethal pediatric solid tumors. High-risk neuroblastoma, characterized by poor prognosis and long-term survival rates below 50% [5, 7], frequently exhibits elevated expression of MYC family oncoproteins, N-MYC and c-MYC [7–9]. These transcription factors are well-known to promote cell proliferation, self-renewal, and an undifferentiated state [10, 11], thereby driving tumor aggressiveness.

Due to their structural properties [12, 13], direct pharmacological targeting of MYC oncoproteins remains largely ineffective and highly challenging, particularly in the context of pediatric cancer. Consequently, alternative therapeutic strategies focus on identifying molecular dependencies imposed by MYC overexpression. In MYC-driven neuroblastoma, mitochondrial gene expression has recently emerged as a promising synthetic lethal target [14]. Elevated MYC levels sensitize tumor cells to mitochondrial ribosome (mitoribosome) inhibitors, which activate the mitochondrial integrated stress response (mitoISR). Induced mitoISR blocks global translation, reducing MYC protein levels and creating synergistic stress that drives selective cell death in MYC-driven tumor cells [14]. However, the molecular mechanisms linking mitoISR activation to MYC-associated tumor cell death remain incompletely understood.

Inducible gene-regulation systems provide a powerful tool for modulating gene expression within the same genetic background, enabling controlled dissection of synthetic lethal interactions, such as those imposed by supraphysiological levels of MYC oncoproteins. Currently, systems relying on tetracycline derivatives as inducers represent one of the most widely used inducible platforms *in vitro* across diverse cell types [15, 16], including MYC-driven neuroblastoma models. However, tetracyclines, as broad-spectrum ribosomal antibiotics [17], exert multiple off-target effects, particularly in the context of mitochondrial biology [18, 19]. Due to the bacterial origin of mitochondria, doxycycline binds to the 28S subunit of the mitoribosome [20], thereby inhibiting mitochondrial translation. As a result, doxycycline disrupts oxidative phosphorylation, induces proteotoxic stress and metabolic reprogramming, and alters mitochondrial morphology, cell proliferation, transcriptional programs, or extracellular matrix remodeling [18–23]. Importantly, such off-target effects occur at concentrations commonly used in tetracycline-inducible systems (5–10 µg/ml after 24 hours and 1 µg/ml after 48 hours [21, 24, 25]), raising concerns that experimental outcomes, particularly in mitochondria-focused studies, may be confounded by the inducer itself [18]. In fact, doxycycline alone currently serves as a tool compound to study mitoribosomal synthetic lethality in MYC-driven neuroblastoma [14].

To overcome these limitations, we sought to establish an alternative MYC-inducible neuroblastoma model using a system that does not interfere with mitochondrial function. We developed novel cumate (p-isopropylbenzoate)-inducible models enabling controlled regulation of c-MYC and N-MYC expression. Although originally described in *Pseudomonas putida* in 1996 [26] and later adapted for prokaryotic and mammalian systems [27, 28], the cumate-regulated gene expression has seen only limited application in human cell models [29–32].

Here, we applied the cumate-inducible system for generating inducer-associated toxicity-free alternatives to widely used tetracycline-based neuroblastoma models, such as Tet21N cells [33]. We demonstrate that cumate does not induce off-target effects on cell viability or mitochondrial function, even at concentrations several-fold higher than those required for effective gene induction. Using this platform, we mechanistically validate MYC-dependent mitoribosomal synthetic lethality and provide the first unbiased evidence that elevated MYC levels sensitize neuroblastoma cells to apoptosis induced by mitochondrial stress. Finally, by reproducing these MYC-dependent synthetic lethal effects in patient-derived cell lines, we identify doxycycline as a strong candidate for repurposing in *MYC(N)*-amplified tumors, highlighting the translational potential of mitoribosomal synthetic lethality for precision therapy in high-risk neuroblastoma.

## 2 RESULTS

### 2.1 Cumate shows no off-target effects on key neuroblastoma cell parameters

To assess cumate (**Fig. 1a**) as a suitable biologically neutral inducer in neuroblastoma models, we first evaluated its effects on cell viability and proliferation. Given our intention to apply the cumate-inducible system to mitochondria-focused research, we also examined the impact of cumate on multiple aspects of mitochondrial function and morphology, as well as on the induction of mitochondrial stress signaling pathways. The tested concentration range of cumate was selected to span the maximum recommended inducer dose according to the manufacturer’s guidelines (50 µg/ml), and effects were assessed after 3-6 days of treatment.

**Figure 1.**
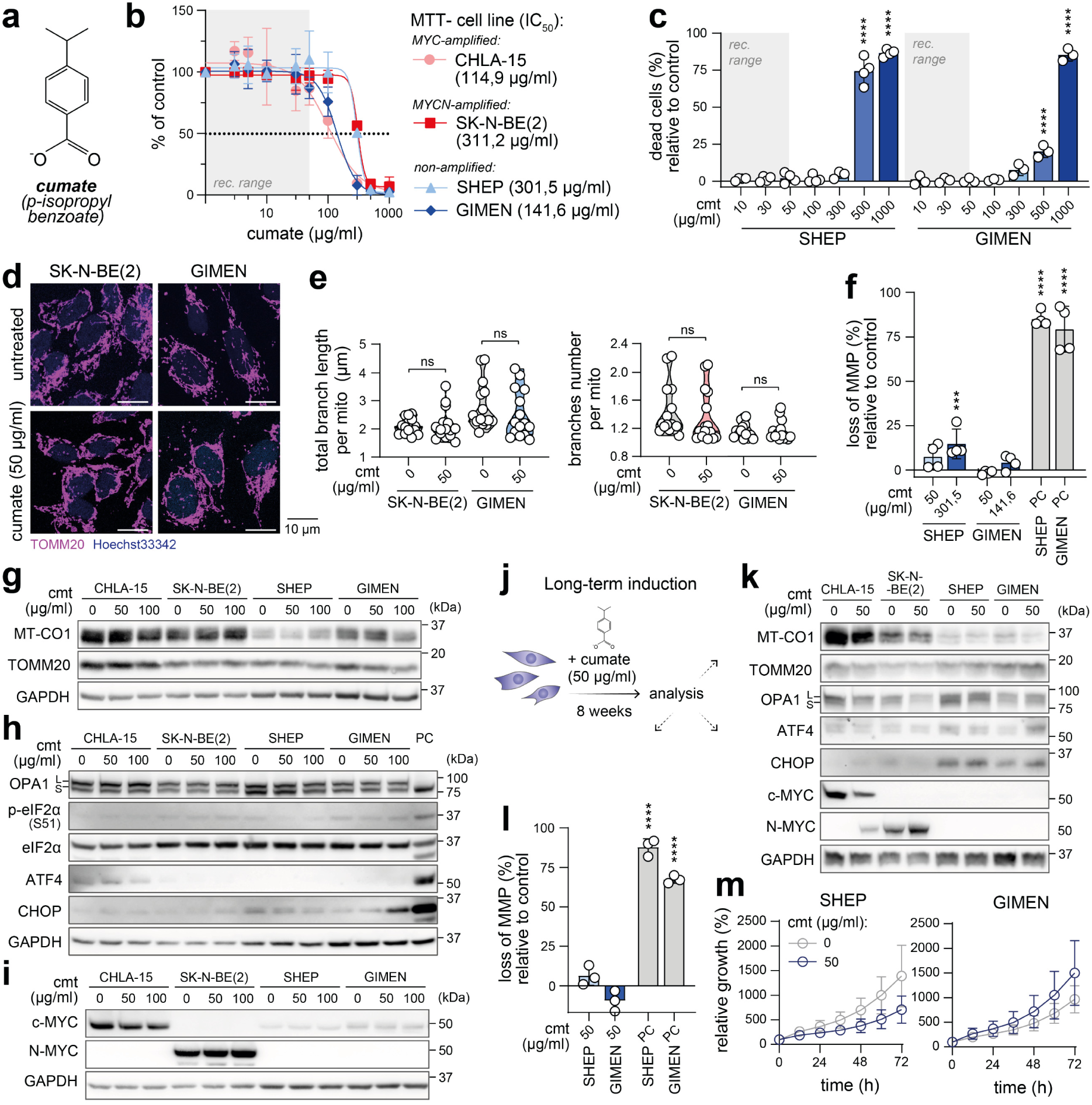
Cumate does not perturb neuroblastoma cells at the maximum recommended dose. (**a**) Molecular structure of cumate (p-isopropyl benzoate). (**b**) Sensitivity of neuroblastoma cell lines to cumate assessed by MTT assay after 72 h of treatment (calculated IC_50_ values are indicated). Data are shown as mean ± SD (biological n = 3, technical n = 3). (**c**) Flow cytometry analysis of cell viability using propidium iodide (PI) staining after 72 h of cumate (cmt) treatment. Data are presented as the difference in percentage of PI-positive cells relative to untreated controls (biological n ≥ 3). (**d**) Mitochondrial morphology visualized by immunofluorescence staining of TOMM20 (magenta) after 72 h treatment with 50 µg/ml cumate. Maximum intensity projections of confocal microscopy Z-stacks are shown. (**e**) Quantitative image analysis using the ImageJ Mitochondria Analyzer plugin. Data represent measurements for individual fields of view (biological n = 3, technical n ≥ 5). (**f**) Flow cytometric analysis of mitochondrial membrane potential (MMP) using the JC-1 probe after 72 h treatment with 50 µg/ml cumate or the respective IC_50_ concentration, relative to untreated controls (biological n = 5). PC, positive control (1h treatment with 10 µM FCCP). (**g-i**) Western blot detection of (**g**) mitochondrial proteins (mitochondrial-encoded MT-CO1 and nuclear-encoded TOMM20), (**h**) markers of mitochondrial stress and ISR, (**i**) c-MYC and N-MYC in a panel of neuroblastoma cells after 72 h cumate treatment. Densitometric analysis is provided in Supplementary Fig. 2. PC, positive control (CHLA-15 cells treated with 50 μM doxycycline for 24 h). (**j-m**) Long-term effects of cumate treatment. (**j**) Schematic representation of the long-term culture experiments. 50 µg/ml cumate was added at each passage for 8 weeks prior to analysis. (**k**) Western blot detection of MT-CO1, markers of ISR and c-MYC in a panel of neuroblastoma cells after 8 weeks of treatment with 50 µg/ml cumate. Densitometric analysis is provided in Supplementary Fig. 2. (**l**) Flow cytometric analysis of MMP after 8 weeks of cumate treatment, relative to untreated controls (biological n = 5). PC, positive control (1h treatment with 10 µM FCCP). (**m**) Live-cell imaging growth rate analysis of SHEP and GIMEN after 8 weeks of cumate treatment. Data are presented as mean ± SD (biological n = 3, technical n = 3). Growth rates of SK-N-BE(2) and CHLA-15 are provided in Supplementary Fig. 6. Statistical significance was determined by one-way ANOVA followed by Dunnett’s multiple comparisons test (**c**,**f**,**l**) and by unpaired t-test with Welch correction (**e,m**), *p < 0.05, **p < 0.01, ***p<0.001, ****p < 0.0001.

At concentrations up to 50 µg/ml, cumate did not affect cell viability and growth across neuroblastoma models (**Fig. 1b; Supplementary Fig. 1a,b**). Cytotoxic or cytostatic effects were observed only at substantially higher, experimentally irrelevant concentrations. Dead-cell staining (**Fig. 1c**), the absence of activated caspase-3, and morphological analysis (**Supplementary Fig. 1c-f**) further excluded apoptosis induction.

50 µg/ml of cumate did not alter mitochondrial morphology (**Fig. 1d,e**), mitochondrial membrane potential (**Fig. 1f**), or mitochondrial protein expression, as assessed by mitochondrially encoded MT-CO1 and nuclear-encoded TOMM20 levels (**Fig. 1g; Supplementary Fig. 2**). In contrast to previously reported effects of doxycycline [14], cumate did not induce the mitochondrial or other forms of integrated stress response (ISR) (**Fig. 1h**), as indicated by unchanged levels of mitochondrial stress sensor OMA1, absence of OPA1 cleavage, and the lack of upregulated eIF2α phosphorylation or ATF4 and CHOP expression. Importantly, cumate treatment also did not affect c-MYC or N-MYC expression levels (**Fig. 1i; Supplementary Fig. 2, 3**). Modest increases in ATF4 and CHOP levels were observed only at substantially higher doses of cumate (**Supplementary Fig. 4, 5**), corresponding to the half-maximal inhibitory concentrations (IC_50_) determined for each cell line (**Fig. 1b**). As these concentrations exceeded the maximum recommended dose by 2–6-fold, mild stress response effects are not unexpected and likely reflect non-specific cellular burden.

Long-term effects were evaluated after 8 weeks of continuous treatment (**Fig. 1j**), during which 50 µg/ml of cumate was added at each passage or medium change to simulate continuous induction. Similarly, no effects were observed on mitochondrial protein expression and stress signaling (**Fig. 1k**), mitochondrial membrane potential (**Fig. 1l**), or cell viability and growth (**Fig. 1m, Supplementary Fig. 6**). Minor changes in MYC protein expression observed in CHLA-15 cells under these conditions were most likely attributable to a phenotypic drift during the long-term culture, as they were not reflected in other examined parameters and detected in other models.

Together, these results demonstrate that cumate provides a well-suited alternative for developing inducible models of MYC-driven neuroblastoma, as it does not significantly affect cell viability, proliferation, or mitochondrial function at concentrations up to 10-fold higher than the effective dose used in our further experiments.

### 2.2 Cumate-inducible neuroblastoma models enable robust and tunable MYC upregulation

Cumate-inducible neuroblastoma models with regulated c-MYC or N-MYC expression were generated by lentiviral transduction using cumate-inducible vectors (**Fig. 2a,b**), containing the *MYC* or *MYCN* coding sequences under the control of a cumate-responsive promoter. The SHEP neuroblastoma cell line, which exhibits low basal expression of MYC family proteins, was used to generate the models. The resulting cell lines, SHEP-CuO-MYC and SHEP-CuO-MYCN, were subsequently characterized for the inducibility of *MYC* and *MYCN* expression, respectively, as well as for the effects of induction on cell proliferation and viability.

**Figure 2.**
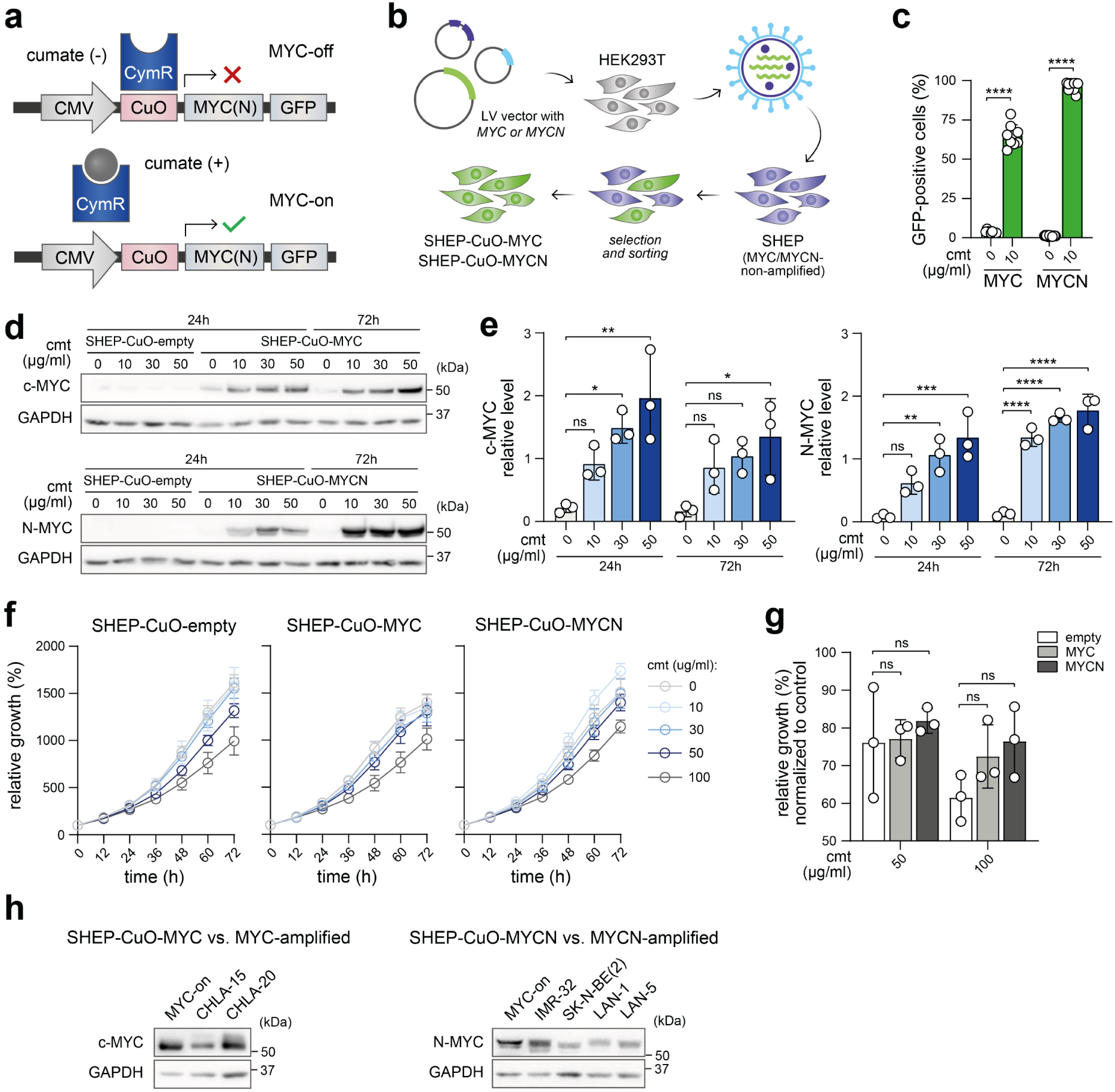
Cumate-inducible models of MYC-driven neuroblastoma. (**a**) Regulation of the cumate-inducible MYC(N). In the absence of cumate, the repressor CymR binds to the CuO operator sequence, preventing transgene expression (MYC-off). When added to the culture media, cumate binds to the repressor, causing its dissociation from the operator and enabling transgene expression (MYC-on). (**b**) Schematic overview of the methodology used to generate stable cumate-inducible neuroblastoma models. (**c**) Flow cytometric analysis of GFP-positive cells 24 h after cumate induction (biological n ≥ 9). (**d**,**e**) Western blot detection (**d**) and densitometric analysis (**e**) of c-MYC and N-MYC protein levels in SHEP-CuO-MYC, SHEP-CuO-MYCN and SHEP-CuO-empty cell lines after cumate induction for 24 h and 72 h. Normalized protein levels are plotted relative to the average of all samples (mean ± SD, biological n = 3). (**f,g**) Live-cell imaging growth rate analysis (**f**) of SHEP-CuO-empty, SHEP-CuO-MYC and SHEP-CuO-MYCN in the presence of indicated concentrations of cumate. (**g**) Confluence after 72h induction with 50 µg/ml and 100 µg/ml of cumate, normalized to untreated controls. Data are presented as mean ± SD (biological n = 3, technical n = 3). (**h**) Levels of c-MYC and N-MYC proteins in SHEP-CuO-MYC and SHEP-CuO-MYCN after induction with 10 µg/ml cumate for 24 h (MYC-on) compared with established *MYC(N)*-amplified neuroblastoma cell lines. Representative western blot (biological n = 3). Statistical significance was determined by one-way ANOVA followed by Dunnett’s multiple comparisons test (**e,g**) and by unpaired two-tailed t-test with Welch correction (**c**), *p < 0.05, **p < 0.01, ****p < 0.0001. cmt, cumate.

The inducibility of the CuO-MYC and CuO-MYCN constructs was validated by an increase in GFP-positive cells (**Fig. 2c**) and elevated c-MYC/N-MYC protein levels (**Fig. 2d,e**), as early as 24 h after addition of cumate to the media. A very low basal c-MYC/N-MYC expression was also detectable in the absence of cumate, indicating partial leakiness of the system. However, even low concentrations of cumate (10 µg/ml) were sufficient to increase MYC expression levels by 4.5- to 8-fold. In addition, the SHEP-CuO-MYC/MYCN models show good reversibility of gene expression, as both c-MYC and N-MYC protein levels return to baseline within 48 h following cumate removal from the cell culture medium (**Supplementary Fig. 7**).

Importantly, growth analysis showed that cumate-induced upregulation of c-MYC and N-MYC levels does not significantly alter cell proliferation compared to the control SHEP-CuO-empty line containing the respective empty vector (**Fig. 2f,g**). These findings indicate that short-term elevation of MYC levels does not affect cellular growth, making it an ideal system for unbiased testing of candidate MYC synthetic lethal drugs. When compared with *MYC(N)*-amplified neuroblastoma cell lines, cumate-induced (MYC-on) SHEP-CuO-MYC/MYCN cells showed similar levels of MYC oncoproteins (**Fig. 2h**), further validating the physiological relevance of the established inducible models.

### 2.3 MYC upregulation primes neuroblastoma cells to doxycycline-induced apoptosis

As a proof-of-concept, we applied the newly established cumate-inducible MYC neuroblastoma models to validate the mitoribosomal synthetic lethality, previously identified by contrasting results in neuroblastoma models with endogenously low or high MYC levels and by using tetracycline-regulated systems [14]. Consistent with earlier observations, doxycycline treatment resulted in significantly increased cell death in cumate-pretreated (MYC-on) cells compared to their uninduced (MYC-off; w/o cumate) counterparts (**Fig. 3a,b**).

**Figure 3.**
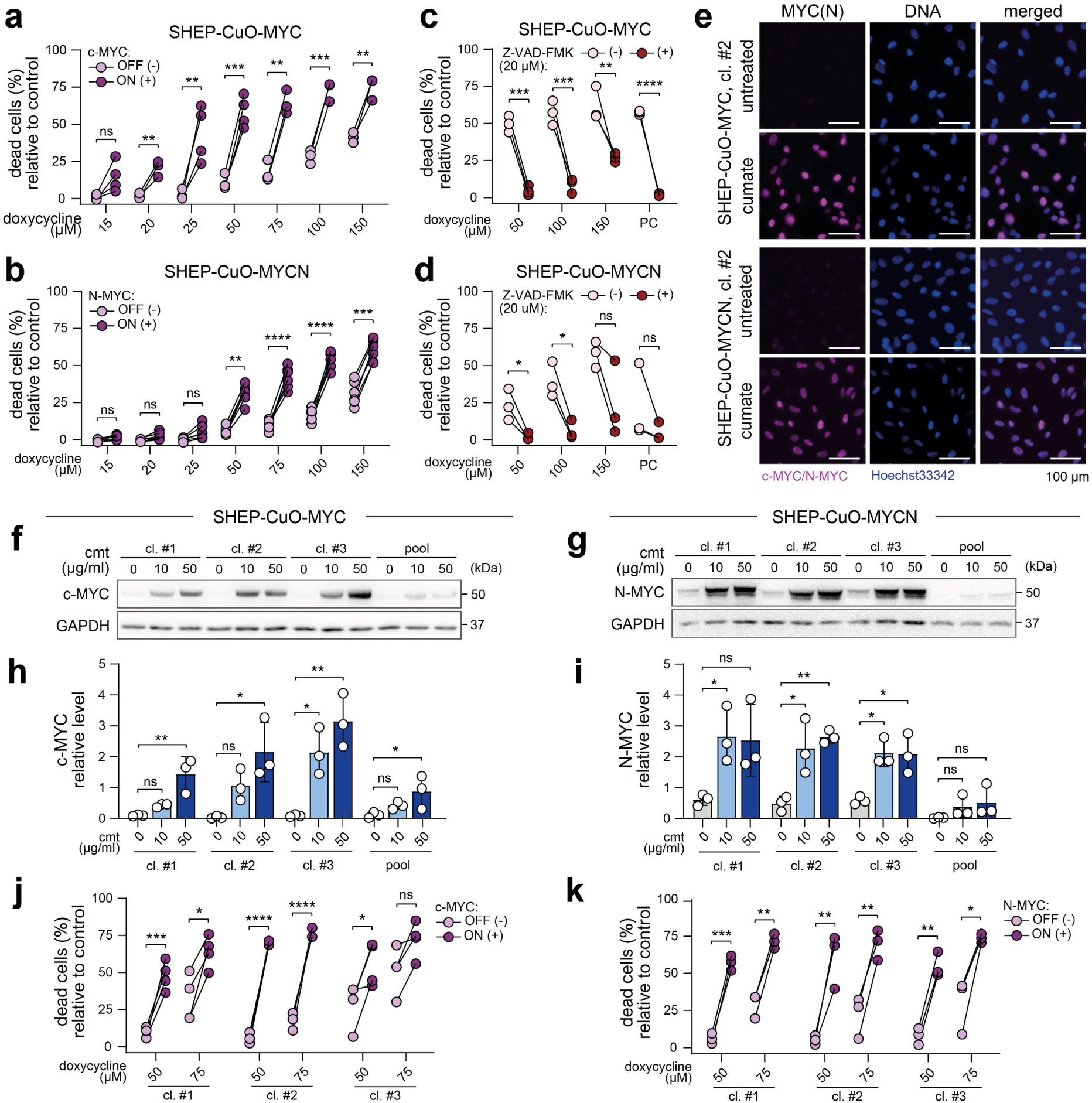
MYC induction enhances neuroblastoma sensitivity to doxycycline-induced apoptosis across both pooled and single-cell clonal backgrounds. **(a, b)** Flow cytometric analysis of cell viability of SHEP-CuO-MYC (a) or SHEP-CuO-MYCN (b) after 24 h of pretreatment with 10 ug/ml cumate (MYC-on) or a vehicle (MYC-off) followed by 24 h doxycycline treatment. Data are presented as the difference in percentage of PI-positive cells relative to doxycycline-untreated controls (biological n ≥ 4). (c, d) Flow cytometric analysis of cell viability of MYC-on SHEP-CuO-MYC (c) or SHEP-CuO-MYCN (**d**) cells (10 µg/ml cumate for 24 h) pretreated with 20 µM Z-VAD-FMK (+) or vehicle (−) for 2 h prior to 24 h doxycycline treatment. Data shown as the difference in percentage of PI-positive cells relative to MYC-off controls (biological n = 3). PC, positive control (24 h treatment with 1µM cisplatin). (**e**) Immunofluorescence staining of c-MYC or N-MYC (magenta) in representative single-cell clones of SHEP-CuO-MYC/MYCN cell lines after 24 h induction with 50 µg/ml cumate. Nuclei were counterstained by Hoechst 33342 (blue). Note the slight leakiness of the inducible system in untreated cells. (**f–i**) Western blot detection (**f, g**) and densitometric analysis (**h,i**) of c-MYC (**f,h**) and N-MYC (**g,i**) protein levels in selected single-cell clones and pool populations of SHEP-CuO-MYC/MYCN cell lines after 24 h of cumate induction. Normalized protein levels are plotted relative to the average of all samples (mean ± SD, biological n = 3). (**j,k**) Flow cytometric analysis of cell viability in MYC-on (10 µg/ml cumate for 24 h) single-cell clones of SHEP-CuO-MYC (**j**) or SHEP-CuO-MYCN (**k**) after 24 h doxycycline treatment. Data are presented as the difference in percentage of PI-positive cells relative to doxycycline-untreated controls (biological n = 3). Statistical significance was determined by one-way ANOVA followed by Dunnett’s multiple comparisons test (**h,i**) and by unpaired two-tailed t-test with Welch correction (**a-d,j,k**), *p < 0.05, **p < 0.01, ***p < 0.001, ****p < 0.0001. cmt, cumate.

Notably, pretreating MYC-on cells with the pan-caspase inhibitor Z-VAD-FMK markedly rescued doxycycline-induced cell death (**Fig. 3c,d**), revealing a regulated caspase-dependent, likely apoptotic mechanism of the MYC synthetic lethality induced by doxycycline-mediated mitoribosome inhibition. The identified caspase-dependent mode of regulated cell death is particularly favorable considering the potential repurposing of doxycycline for the treatment of high-risk MYC-driven neuroblastoma.

To account for cellular heterogeneity that might contribute to the observed synthetic lethal effects, we next sought to determine whether MYC-dependent selectivity is conserved at the level of individual cell clones. Single-cell clones exhibiting the highest GFP signal upon cumate treatment (marking efficient inducibility of CuO-MYC and CuO-MYCN constructs) were isolated by FACS and further characterized by immunodetection. While MYC expression levels still varied among individual cells (**Fig. 3e**), selected single-cell clones displayed markedly higher MYC levels upon cumate induction than the parental pool populations (**Fig. 3f–i**). Importantly, doxycycline-induced apoptosis was preserved in all three tested clones of both SHEP-CuO-MYC and SHEP-CuO-MYCN cells (**Fig. 3j,k**), demonstrating that the sensitivity does not arise from averaging responses across population-level heterogeneity.

### 2.4 MYC degradation, not altered mitochondrial stress response, marks mitoribosomal synthetic lethality in MYC-driven cells

To elucidate the mechanisms underlying the increased sensitivity of MYC-on cells to doxycycline, we analyzed mitochondrial stress signaling and mitochondrial morphology following doxycycline treatment. Despite ectopic MYC overexpression, doxycycline caused a marked, dose-dependent reduction in c-MYC/N-MYC protein levels in MYC-on cells (**Fig. 4a-d**), consistent with the previously described mechanism of action that favors rapid proteasomal degradation of MYC oncoproteins due to the mitoISR-inhibited translation of MYC transcripts [14]. Indeed, doxycycline selectively impaired mitochondrial translation, documented by the reduced levels of mitochondrially encoded MT-CO1 (**Fig. 4e**). Analysis of mitoISR components further demonstrated that the doxycycline-induced mitochondrial proteotoxic stress resulted in activation of the OMA1 protease, as marked by its autolytic degradation and the reduction of the long isoform of OPA1, a known OMA1 substrate (**Fig. 4e**). In line with previous studies [14, 34, 35], the stress signaling was relayed to the cytosolic components of the ISR, manifested by increased phosphorylation of eIF2α and upregulation of ATF4 and CHOP levels in a concentration-dependent manner (**Fig. 4e**).

**Figure 4.**
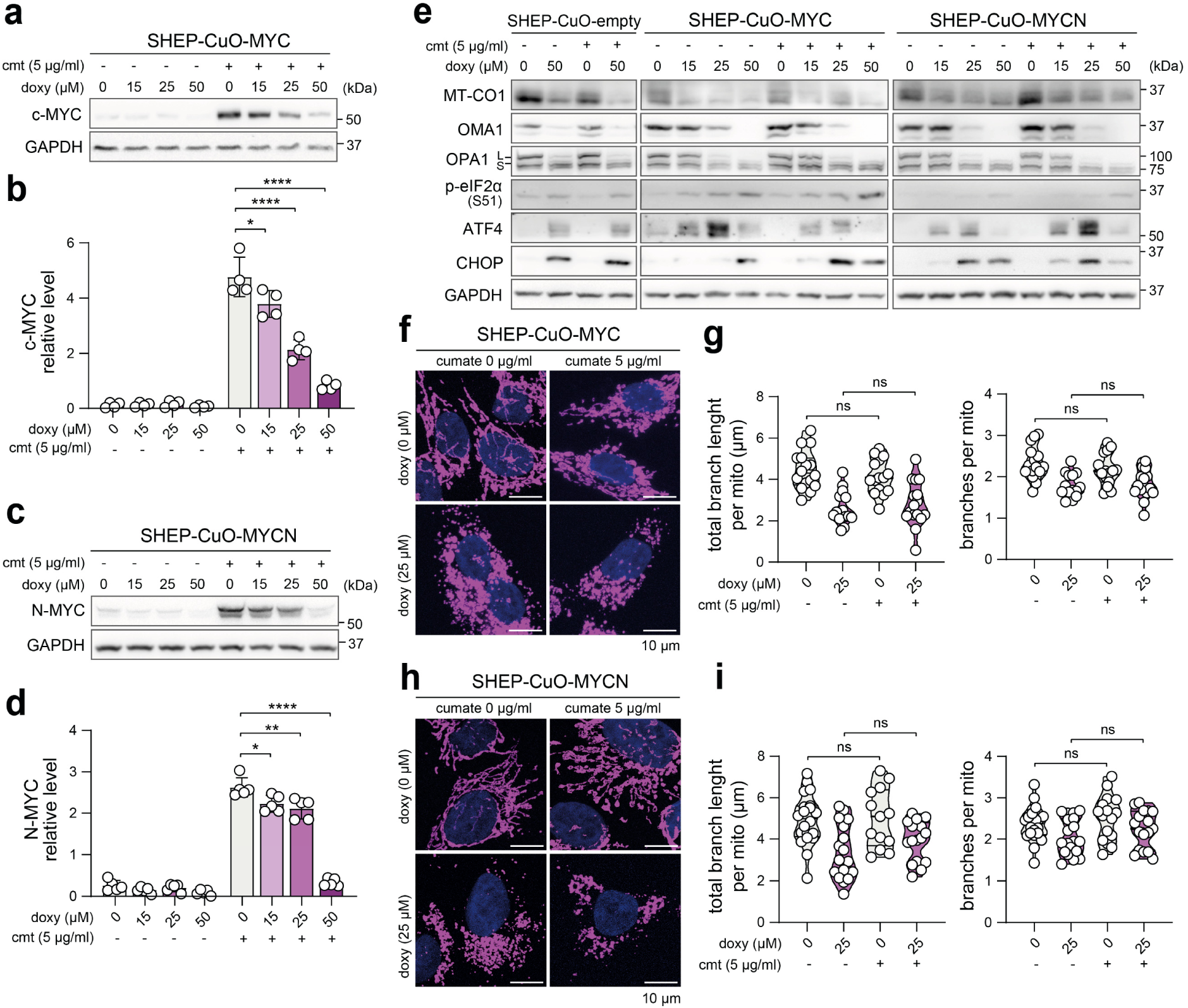
Doxycycline induces a uniform, MYC-independent mitoISR selectively coupled to strong MYC downregulation in MYC-overexpressing cells. **(a-d)** Western blot detection (a,c) and densitometric analysis (b,d) of c-MYC and N-MYC protein levels, respectively, across SHEP-CuO-MYC/MYCN single-cell clones after cumate induction for 24 h followed by 24 h of doxycycline treatment. Normalized protein levels are plotted relative to the average of all samples (mean ± SD, biological n ≥ 4). (e) Western blot detection of MT-CO1 and markers of mitoISR in selected SHEP-CuO-empty, SHEP-CuO-MYC, and SHEP-CuO-MYCN single-cell clones after 24 h of cumate induction followed by doxycycline treatment for 24 h. Densitometric analysis is provided in Supplementary Fig. 8. (f-i) Mitochondrial morphology visualized by immunofluorescence staining of TOMM20 (magenta) (f,h) in selected single-cell clone-derived SHEP-CuO-MYC/MYCN cell lines after 24 h induction followed by 24 h of treatment with 25 µM doxycycline. Maximum intensity projections of confocal microscopy Z-stacks are shown. Quantitative image analysis (g,i) using the ImageJ Mitochondria Analyzer plugin. Data are presented as parameters determined for individual field of vision images (biological n = 3, technical n ≥ 4). Statistical significance was determined by one-way ANOVA followed by Dunnett’s multiple comparisons test (b,d) and by unpaired two-tailed t-test with Welch correction (g,i), *p < 0.05, **p < 0.01, ****p < 0.0001. cmt, cumate; doxy, doxycycline.

Importantly, the degree of mitoISR activation did not differ between MYC-on and MYC-off states (**Supplementary Fig. 8**), suggesting that doxycycline-induced mitoISR is independent of MYC expression. This was further confirmed in control SHEP-CuO-empty cells carrying the empty vector, which showed a similar level of mitoISR activation in response to doxycycline treatment (**Fig. 4e; Supplementary Fig. 8**). Thus, the increased sensitivity of MYC-on cells to doxycycline-induced cell death cannot be attributed to enhanced mitochondrial stress signaling. Instead, the critical distinction appears to be the rapid degradation of MYC proteins following doxycycline treatment in MYC-overexpressing cells (**Fig. 4a-d**). It remains unclear whether this dramatic downregulation contributes to cell death through disrupting cellular programs imposed by elevated MYC levels or through other mechanisms. However, our results show that, in a genetically uniform background, MYC overexpression prior to mitoribosome inhibition is a strong predictor of cell death induction. By analyzing GFP signal intensity, corresponding to MYC levels, and the percentage of dead cells across both pool populations and selected single-cell clones, we confirmed that cells with higher MYC levels are more susceptible to doxycycline-induced apoptosis (**Supplementary Fig. 9**). This correlation was particularly pronounced in SHEP-CuO-MYC cells. In SHEP-CuO-MYCN, this effect was likely masked due to the uniformly high percentage of GFP-positive (MYCN-overexpressing) cells across the analyzed clones and pool populations.

Consistent with the mitoISR activation, analysis of mitochondrial morphology revealed a similar increase in stress-induced mitochondrial fragmentation following doxycycline treatment in both MYC-off and MYC-on conditions (**Fig. 4f-i**). These results demonstrate that doxycycline-induced mitochondrial stress occurs independently of MYC levels and does not mark cells sensitive to the mitoribosome inhibition. The mitoISR has previously been shown to block global protein synthesis and to shift MYC protein turnover toward proteasomal degradation [14]. Our mechanistic findings support these observations, suggesting that coupling of mitochondrial stress to the rapid loss of MYC oncoproteins, rather than the stress alone, contributes to apoptotic cell death in MYC-on cells adapted to high MYC levels.

### 2.5 Cumate-inducible MYC models generalize mitoribosomal synthetic lethality and reveal context-dependent activity of ribosomal antibiotics

We next sought to assess the utility of cumate-inducible models as a platform for screening and profiling the effects of other bacterial ribosome-targeting antibiotics. Two well-characterized drugs, tigecycline and chloramphenicol, with different primary target sites at the bacterial ribosome and corresponding mitoribosome subunits [12, 36, 37] were used for these proof-of-concept experiments.

Consistent with doxycycline, tigecycline, another tetracycline-derived antibiotic, induced higher levels of cell death in MYC-on cells than in MYC-off conditions (**Fig. 5a**). However, the magnitude of the effect was weaker than with doxycycline. In SHEP-CuO-MYC cells, treatment of MYC-on cells with 100µM tigecycline for 24 h resulted in approximately 30-40% cell death, compared to approximately 75% following treatment with doxycycline at the same concentration (**Fig. 3a**). Nevertheless, the relative increase in cell death between MYC-on and MYC-off conditions was comparable for both drugs. N-MYC-driven SHEP-CuO-MYCN cells were overall less sensitive, yet tigecycline treatment resulted in significantly increased cell death under MYC-on conditions (**Fig. 5a**).

**Figure 5.**
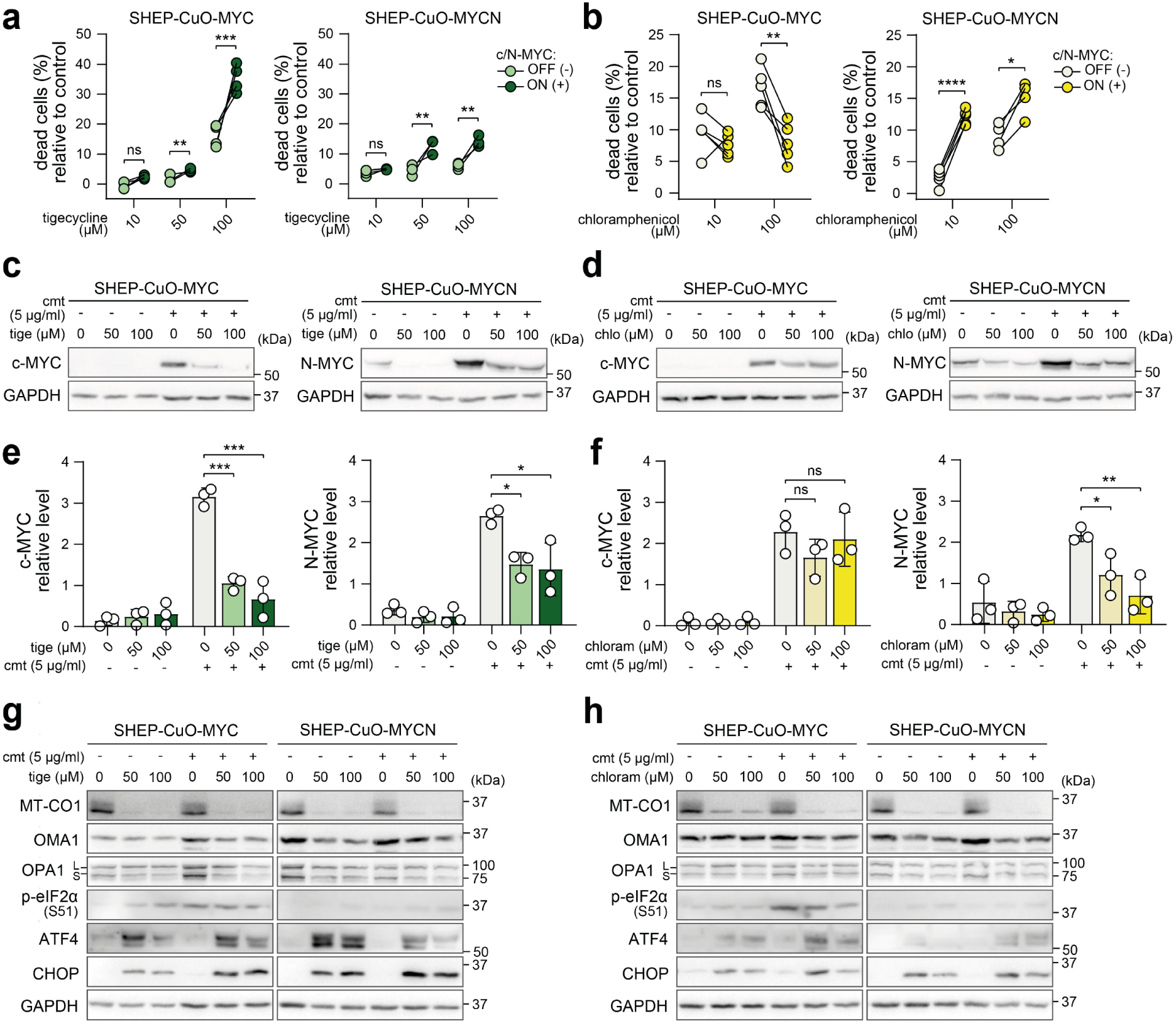
MYC synthetic lethality of mitoribosome-inhibiting antibiotics converges on coupling mitochondrial stress-activated ISR with efficient MYC degradation. **(a,b)** Flow cytometry analysis of cell viability using PI staining in SHEP-CuO-MYC or SHEP-CuO-MYCN after at least 24 h of treatment with 5 µg/ml cumate (MYC-on) followed by 24 h of treatment with tigecycline (a) or chloramphenicol (b). Data are presented as the difference relative to antibiotic-untreated controls (biological n = 3). (c-f) Western blot detection (c,d) and densitometric analysis (e,f) of c-MYC and N-MYC protein levels in selected single-cell clone-derived SHEP-CuO-MYC/MYCN cell lines after at least 24 h of treatment with 5 µg/ml cumate followed by 24h of treatment with indicated tigecycline (c,e) or chloramphenicol (d,f) concentrations. Normalized protein levels are plotted relative to the average of all samples (mean ± SD, biological n ≥ 4). (g,h) Western blot detection of MT-CO1 and markers of ISR in selected single-cell clone-derived SHEP-CuO-MYC/MYCN cell lines after at least 24 h of treatment with 5 µg/ml cumate followed by 24 h of treatment with indicated tigecycline (g) or chloramphenicol (h) concentrations. Densitometric analysis is provided in Supplementary Fig. 10. Statistical significance was determined by one-way ANOVA followed by Dunnett’s multiple comparisons test (b) and by unpaired two-tailed t-test with Welch correction (f,h), *p < 0.05, **p < 0.01, ****p < 0.0001. chloram, chloramphenicol; cmt, cumate; tige, tigecycline.

Unexpectedly, inducing c-MYC expression in SHEP-CuO-MYC cells led to reduced cell death in response to chloramphenicol treatment (**Fig. 5b**). In contrast, SHEP-CuO-MYCN cells showed the expected enhanced sensitivity to chloramphenicol in the MYC-on state. Mechanistically, tigecycline downregulated c-MYC and N-MYC protein levels (**Fig. 5d,e**), which was consistent with doxycycline effects. Impaired mitochondrial translation was confirmed by reduced MT-CO1 expression, and tigecycline-induced mitoISR activation was comparable between MYC-on and MYC-off states (**Fig. 5h; Supplementary Fig. 10a,b**). In contrast, although chloramphenicol activated mitoISR (**Fig. 5i**), it led to significantly reduced levels of N-MYC but not c-MYC (**Fig. 5g**), which may potentially explain the absence of enhanced cell death in MYC-on SHEP-CuO-MYC cells.

In summary, tigecycline phenocopies doxycycline effects by preferentially inducing cell death in MYC-overexpressing cells, while chloramphenicol shows context-dependent activity. These results demonstrate the utility of our cumate-inducible neuroblastoma models for identifying mitochondrial MYC synthetic lethal interactions and exploring factors that may fine-tune responses to mitochondrial stress signaling. Our findings provide orthogonal evidence supporting previous observations [14] and indicate that rapid degradation of MYC oncoproteins following mitoISR activation by mitoribosomal inhibitors is a key mechanism underlying synthetic lethal effects in MYC-driven neuroblastoma.

### 2.6 Doxycycline recapitulates MYC-synthetic lethal effects across clinically relevant patient-derived neuroblastoma cell lines

Finally, to evaluate the translational relevance of findings obtained in our cumate-inducible models, we tested doxycycline in three low-passage neuroblastoma cell lines derived in-house from patient tumor specimens: one *MYCN*-amplified (NBL-58) and two lacking *MYC(N)* amplification (NBL-28 and NBL-38). Responses were compared with those observed in established *MYC(N)*-amplified neuroblastoma models.

Consistent with previous findings, the *MYCN*-amplified NBL-58 cells were significantly more sensitive to doxycycline-induced cell death than their non-amplified counterparts, NBL-38 and NBL-28 (**Fig. 6a**). While activation of the mitoISR was conserved across all models, marked reduction of MYCN levels was observed in NBL-58 cells (**Fig. 6b-d**), reflecting comparable MYC-selective responses among patient-derived and established neuroblastoma cell lines.

**Figure 6.**
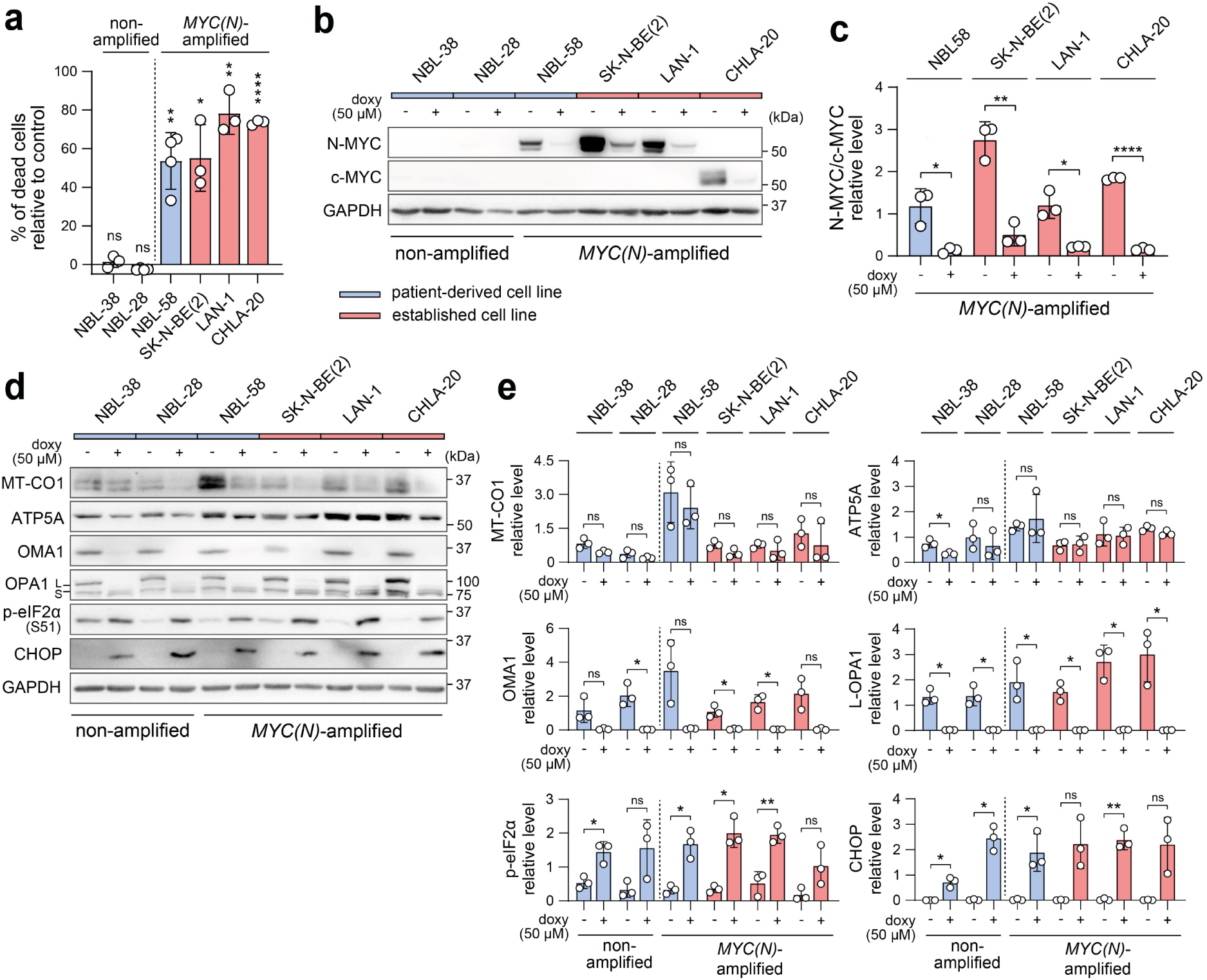
Patient-derived neuroblastoma cell lines validate the translational potential of doxycycline-induced MYC synthetic lethality. **(a)** Flow cytometry analysis of cell viability using PI staining in patient-derived cell lines (NBL-38, NBL-28, NBL-58) and established cell lines (SK-N-BE(2), LAN-1, CHLA-20) after 48 h of treatment with 100 µM doxycycline. Data are presented as the difference in percentage of PI-positive cells relative to doxycycline-untreated controls (biological n = 3–4). (b,c) Western blot detection (b) and densitometric analysis (c) of N-MYC or c-MYC protein downregulation in patient-derived and established cell lines after 24 h of treatment with 50 µM doxycycline. Normalized protein levels are plotted relative to the average of all samples (mean ± SD, biological n = 3). (d,e) Western blot detection (d) and densitometric analysis (e) of MT-CO1, ATP5A and mitoISR markers after 24h of treatment with 50 µM doxycycline. Note the activation of mitoISR upon doxycycline treatment conserved across all models. Normalized protein levels are plotted relative to the average of all samples (mean ± SD, biological n = 3). Statistical significance was determined by unpaired two-tailed t-test with Welch correction (a,c,e), *p < 0.05, **p < 0.01, ***p < 0.001, ****p < 0.0001. doxy; doxycycline.

Together, these findings demonstrate that the MYC-synthetic lethal effects of doxycycline are not restricted to established neuroblastoma models. Their reproducibility in low-passage patient-derived neuroblastoma cell lines supports the clinical relevance of mitoribosomal synthetic lethality and further validates the predictive value of our cumate-inducible MYC models.

## 3 DISCUSSION

MYC overexpression imposes distinct dependencies, creating attractive opportunities for synthetic lethal targeting in MYC-driven cancers [38]. However, accurate interrogation of these dependencies requires molecular tools that modulate MYC activity without confounding effects on cellular physiology. Here, we established novel cumate-inducible neuroblastoma models that enable controlled *MYC(N)* expression without the mitochondrial or cytotoxic off-target effects associated with commonly used inducers, providing a reliable platform for functional studies of MYC biology in neuroblastoma.

Given the relatively recent introduction of cumate-regulated systems and their limited use in mammalian cells [27–31], validation of cumate’s biological inertness has been lacking. We demonstrate that cumate does not affect key cellular parameters, including mitochondrial function, stress signaling, or c-MYC/N-MYC levels, even at the highest concentrations recommended for gene induction. This addresses a major limitation of tetracycline-inducible systems, which rely on inducers known to elicit mitochondrial and proteostatic stress at commonly used doses [18, 21, 23, 25]. Collectively, these findings establish cumate-regulated systems as a more suitable alternative for synthetic lethal studies that require precise control of MYC, or other target genes, within a uniform genetic background.

Using SHEP cells with very low endogenous MYC levels, we generated cumate-inducible MYC neuroblastoma models that enable biologically relevant c-MYC or N-MYC overexpression. Despite low-level leakiness, consistent with previous reports using cumate-regulated systems [29, 30, 32, 39], cumate induction produced a clear separation between MYC-on and MYC-off states. Both proteins significantly increased within 24 h of induction with 10 µg/ml cumate and progressively accumulated toward steady-state levels during prolonged exposure (Fig. 2d,e). Notably, even 5 µg/ml of cumate was sufficient to markedly elevate MYC protein levels (Fig. 4) and unmask synthetic lethal effects of mitoribosome inhibition in MYC-on cells. Although higher concentrations of cumate (>10 µg/ml) retained dose-dependent activity, confirming that the system is titratable, they are unlikely to yield additional biologically and experimentally meaningful benefits.

Although the dynamic range of cumate-inducible expression is lower than that of some tetracycline-based platforms [15, 40, 41], induced MYC levels closely recapitulated those detected in *MYC(N)*-amplified neuroblastoma cells. Such clinically relevant inducibility is ideal for most applications, particularly for testing drug candidates, which underpins the physiological relevance of the established SHEP-CuO-MYC/MYCN models. When required, next iterations could potentially expand the dynamic range via a repressible cumate-switch configuration, analogous to the Tet-Off system used in the Tet21N neuroblastoma model [14, 42]. Additionally, as previously reported, MYC accumulation may eventually reach a threshold that triggers cell death [43–45]. This is an inherent constraint limiting some long-term applications of our models. Despite the neuroblastoma cell context, known to better tolerate elevated MYC levels, sustained supraphysiological MYC induction for more than three weeks markedly reduced cell viability, precluding further analyses. However, as 8-week exposure to the highest recommended cumate dose did not impair viability of neuroblastoma cells lacking the cumate-inducible *MYC(N)* constructs (Fig. 1j-k), the toxicity observed in MYC-on cells can be fully attributed to long-term MYC induction, likely activating MYC-driven pro-apoptotic pathways.

In contrast, we did not observe compromised cell viability after mid-term induction (approximately three weeks/10 passages), indicating that MYC-mediated cell death is both time- and level-dependent. Notably, even moderate MYC upregulation significantly altered sensitivity to mitoribosome inhibition, suggesting that the system recapitulates, at least in part, physiologically relevant MYC-driven programs observed in *MYC(N)*-amplified neuroblastoma [14]. Comparison with empty vector controls showed that the canonical growth-promoting effect of MYC oncoproteins [11, 46, 47] was not significant within the experimental timeframe. This excludes differential proliferation as a confounding factor and enables mapping of drug responses to MYC-dependent programs that extend beyond proliferation.

To demonstrate the utility of cumate-inducible MYC models, we performed proof-of-concept experiments focusing on mitochondrial MYC-dependent synthetic lethal effects of ribosomal antibiotics. Corroborating findings of our previous study [14], we provide unbiased mechanistic evidence that elevated MYC levels sensitize neuroblastoma cells to doxycycline, a ribosomal antibiotic targeting both bacterial and mitochondrial protein synthesis [20]. Crucially, we show for the first time that the resulting cell death is mediated by a caspase-dependent apoptotic program that can be rescued by pan-caspase inhibition. A similar, albeit less pronounced, effect was observed following treatment with tigecycline and chloramphenicol (Fig. 5), which likewise inhibit mitochondrial ribosomes [48, 49]. These results indicate that MYC synthetic lethality is broadly associated with mitoribosome inhibition and can be readily captured by our experimental platform.

Mechanistically, inhibition of mitochondrial translation induced mitochondrial stress and activated mitoISR in both MYC-off and MYC-on cells (Fig. 4e-i). However, only MYC-on cells exhibited rapid MYC downregulation (Fig. 4a-d) and subsequent apoptotic cell death, indicating a specific vulnerability of MYC-overexpressing cells to mitoribosome inhibition. Notably, this MYC-imposed vulnerability to doxycycline-induced mitoISR was evident at concentrations within the clinically safe range used in children [50–52]. Importantly, our study further extends the concept of MYC-dependent mitoribosomal synthetic lethality by demonstrating its reproducibility in low-passage patient-derived neuroblastoma cells, supporting the clinical relevance of mitochondrial translation inhibition as a potential precision oncology strategy for MYC-driven neuroblastoma. Together with previous observations [14], these findings identify doxycycline as a compelling candidate for drug repurposing and provide a strong rationale for its further preclinical and clinical evaluation in *MYC(N)*-amplified neuroblastoma.

More broadly, the capacity of our SHEP-CuO-MYC/MYCN models to reveal clinically relevant MYC-dependent drug responses highlights their value as a scalable platform for screening FDA-approved compounds to identify additional synthetic-lethal hits. Such drug-repurposing applications are particularly attractive in pediatric oncology, where therapeutic options remain largely limited to modified dosing of anticancer drugs originally developed for adults, with only a few specifically designed for pediatric malignancies [53–58]. Given its extensive clinical use and favorable safety profile, doxycycline represents a promising first example of how mitochondria-perturbing drugs could be repurposed for *MYC(N)*-amplified neuroblastoma. Future *in vivo* studies will be required to define the therapeutic window of this approach and to determine whether mitochondrial MYC-dependent synthetic lethality can be translated into a broader precision medicine strategy for patients with high-risk neuroblastoma, and potentially other MYC-driven cancers.

## 4 METHODS

### 4.1 Ethics

The study was conducted in accordance with the Declaration of Helsinki and approved by the Masaryk University Research Ethics Committee (ref. no. EKV-2024-031; granted on December 9, 2024) and the Ethics Board of the Faculty of Medicine (ref. no. 0299/2025; granted on March 14, 2025). All noncommercial cell lines were derived from tumor tissues with written informed consent obtained from patients or their legal guardians.

### 4.2 Cell culture and treatment

The following established cell lines were used in this study: (i) neuroblastoma cell lines CHLA-15 and CHLA-20 (Cellosaurus #CVCL_6594 and #CVCL_6602, kindly provided by Prof. Michael Hogarty, Children’s Hospital of Philadelphia, USA), SHEP (#CVCL_0524, generously provided by Dr. Frank Westermann, DKFZ, Heidelberg, Germany), GIMEN (#CVCL_1232), IMR-32 (#CVCL_0346), LAN-1 (#CVCL_1827) LAN-5 (#CVCL_0389; all kindly provided by Prof. Lumír Krejčí, Masaryk University, Brno), SK-N-BE(2) (#CVCL_0528, purchased from ECACC), (ii) human embryonic kidney cell line HEK293T (#CVCL_0063, a kind gift of Dr. Michal Šmída, CEITEC, Masaryk University, Brno). Patient-derived primary neuroblastoma cell lines NBL-28, NBL-38, and NBL-58 were established from tumor tissue samples as described elsewhere [59]. Written informed consents were obtained under research projects (IGA MZCR NR/9125-4, AZV MZCR 15-34621A, and University Hospital Brno 14/24) approved by the respective research ethics bodies of Masaryk University and University Hospital Brno (approval no. 23/2005, 26/08/2014, and 06-140224/EK, respectively). All cell lines were authenticated by STR profiling (Generi Biotech, Hradec Králové, Czech Republic) and routinely tested for mycoplasma contamination via PCR [60]. The composition of culture media used is described in Supplementary Table 1. All cell lines were cultured at 37 °C in a humidified atmosphere containing 5% CO_2_.

Drug treatments were always performed the day after cell seeding. The following drugs, inducers or inhibitors were used for treatment: cumate (#QM150A-1, System Biosciences, Palo Alto, California, USA), doxycycline hyclate (#D9891, Sigma-Aldrich, St. Louis, Missouri, USA), tigecycline hydrate (#PZ0021, Sigma-Aldrich), chloramphenicol (#1678502, Serva, Heidelberg, Germany), Z-VAD-FMK (#HY-16658, MedChemExpress, New Jersey, USA), cisplatin (#232120, Sigma-Aldrich, St. Louis, Missouri, USA).

### 4.3 Cumate-inducible MYC and MYCN vectors

MYC sequence was subcloned and amplified by cloning PCR from the donor plasmid pcDNA3.3_c-MYC [61] kindly provided by Derrick Rossi (Addgene plasmid #26818; http://n2t.net/addgene:26818; RRID:Addgene_26818). The primer sequences used are listed in Supplementary Table 3. The fragment was separated on 1% agarose gel and isolated using NucleoSpin Gen and PCR Clean-up (#740609.50, Macherey-Nagel, Düren, Germany). The MYCN sequence with respective restriction sites was prepared by gene synthesis (#CLPMX5UGDE, Invitrogen) and excised from the original pMX vector backbone using restriction endonucleases. MYC or MYCN sequence was ligated into cumate-inducible pCDH-CuO-MCS-IRES-GFP-EF1α-CymR-T2A-Puro SparQ™ (pCDH-CuO) lentiviral vector (QM812B-1, System Biosciences). For plasmid amplification, *E. coli* Stbl3 cells were transformed with the pCDH-CuO vector coding for MYC (pCDH-CuO-MYC) or MYCN (pCDH-CuO-MYCN) gene under cumate-inducible promoter, plated on LB-agar with 100 µg/ml ampicillin, and grown overnight at 37 °C. The next day, a single colony was transferred into liquid LB media supplemented with 100 µg/ml ampicillin and incubated on a roller overnight (200 rpm, 37°C). Plasmid isolation was performed using NucleoSpin Plasmid kit (#740588.10, Macherey-Nagel). The cloning efficiency was verified by whole-plasmid sequencing (Eurofins Genomics, Brno, Czech Republic).

### 4.4 Generation of cumate-inducible SHEP-CuO-MYC/MYCN cell lines and single-cell clones

Lentiviral particles were generated as described previously [62, 63]. Briefly, HEK293T cells were transfected with pCDH-CuO empty vector (CuO-empty) or one of the generated pCDH-CuO-MYC or pCDH-CuO-MYCN vectors together with the 2^nd^ generation of lentiviral production plasmids psPAX2 (#12260, Addgene) and pMD2.G (#12259, Addgene), kindly provided by Didier Trono, all mixed with PEI (#HY-K2014, MedChemExpress). After transfection, the cell culture medium was exchanged for OptiMEM (#31985062, Invitrogen) containing 1% FBS, 1% MEM non-essential amino acid solution, and 1% penicillin/streptomycin. The medium was collected every 12 h for a total of 48 h and kept at 4°C. The virus supernatant was centrifuged (4500 × g, 5 min, room temperature) and filtered through a 0.45 µm low-protein-binding filter.

The virus supernatant was mixed with Polybrene (#28728-55-4, Santa Cruz Biotechnology) at a final concentration of 10 μg/ml and applied to SHEP cells overnight. The next day, the medium containing viral particles was replaced with a fresh culture medium. Transduced cells were then continuously cultured in the presence of 2.5 µg/ml puromycin (#sc-108071, Santa Cruz Biotechnology) to select and maintain stable SHEP-CuO-MYC, SHEP-CuO-MYCN, and SHEP-CuO-empty cell lines. To induce MYC or MYCN expression (MYC-on), cells were cultured in the presence of cumate for at least 24 h prior to analysis or drug treatments. SHEP-CuO-MYC/MYCN cells cultured in the absence of cumate (MYC-off) or SHEP-CuO-empty cells were used as controls.

Single-cell clones were generated from SHEP-CuO-MYC and SHEP-CuO-MYCN cell pools by fluorescence-assisted cell sorting based on GFP expression following cumate induction, using BD FACSAria Fusion flow cytometer (BD Biosciences, Waters Corporation, Heidelberg, Germany).

### 4.5 Growth analysis and MTT assay

Cells were seeded into 96-well plates (CHLA-15: 5000 cells/well for 72h, 2500 cells/well for 6d; SK-N-BE(2): 2000 cells/well for 72h, 1000 cells/well for 6d; SHEP, GIMEN: 1000 cells/well for 72h, 750 cells/well for 6d) and treated the day after seeding. Cell confluency was determined every 4 or 6h for 72h or 6d using a live-cell imaging system Incucyte® SX1 (Sartorius, Göttingen, Germany) and plotted relative to initial values. For MTT assay, after the incubation time, thiazolyl blue tetrazolium bromide (#M2128, Sigma-Aldrich) was added to each well to reach the final concentration of 0,455 mg/ml. After 3h incubation under standard conditions, the medium was replaced with 200 μl of DMSO to solubilize the formazan crystals. The absorbance of each well was determined by Sunrise Absorbance Reader (Tecan, Männedorf, Switzerland). The half-maximal inhibitory concentrations (IC_50_) of drugs were determined via nonlinear regression of the MTT assay data using GraphPad Prism 8.0.2 software (GraphPad Software, San Diego, California, USA), and the absolute IC_50_ values were determined according to the following formula: relative IC_50_*(((50-top)/(bottom-50))^(− 1/hill slope)).

### 4.6 Western blotting

Lysates were collected using RIPA lysis buffer (2 mM EDTA, 1% IGEPAL® CA-630, 0.1% SDS, 8.7 mg/ml sodium chloride, 5 mg/ml sodium deoxycholate, 50 mM Tris-HCl) supplemented with cOmplete™ Mini Protease Inhibitor Cocktail (#11836170001, Roche, Basel, Switzerland) and PhosSTOP (#4906837001, Roche). The samples were resolved by SDS-PAGE on 10% polyacrylamide gels and transferred onto PVDF membranes (#1620177, Bio-Rad Laboratories, Hercules, CA, USA). Blocking was performed in 5% not-fat dry milk or bovine serum albumin (#A7906, Sigma-Aldrich) in Tris-buffered saline with 0.05% Tween-20 (#93773, Sigma-Aldrich) for at least 1 h, followed by incubation with primary antibodies overnight on a rocking platform at 4 °C and by the incubation with secondary HRP-linked antibody at RT for at least 1 h. The list of antibodies used, including dilutions and respective blocking agents, is provided in Supplementary Table 2. Chemiluminescence was detected after a 5-min incubation with ECL™ Prime Western Blotting Detection Reagent (#RPN2236, Cytiva, Marlborough, Massachusetts, USA) using Azure C600 imaging system (Azure Biosystems, Dublin, California, USA). Densitometric analysis of western blotting images was performed using the gel analysis tool in ImageJ (Fiji) software (NIH, Bethesda, MD, USA), version 2.1.0/1.53c. The signal of a protein of interest was normalized to that of a loading control, α-tubulin or GAPDH, detected on the same gel.

### 4.7 Immunostaining

Cells were seeded on coverslips coated with Matrigel (#734-1440, Corning, New York, USA) and treated with respective drugs. After the incubation period, the coverslips were rinsed by PBS, cells were fixed by 3% paraformaldehyde (#158127, Sigma-Aldrich) and permeabilized using 0.2% Triton X-100 (#04807423, MP Biomedicals, Irvine, California, USA) for 1 min. Blocking was performed by 3% bovine serum albumin (#A7906, Sigma-Aldrich) for 10 min at RT. The primary and secondary antibodies (Supplementary Table 2) were diluted in blocking solution. The incubation with antibodies lasted for at least 60 min. Nuclei were stained by Hoechst 33342 (Life Technologies). The cells were mounted with ProLong™ Diamond Antifade (#P36961, Invitrogen, Carlsbad, California, USA) and imaged using Zeiss Axio Imager Z2 microscope (Zeiss, Jena, Germany). For mitochondria morphology analysis, cells were imaged using Leica SP8 confocal microscope (Leica, Wetzlar, Germany). Z-stacked images were captured and processed as maximum intensity projections using LAS X software (Leica, 3.4.218368).

### 4.8 Mitochondrial morphology analysis

Mitochondrial network morphology was analyzed on a per-field-of-view basis from 2D maximum intensity projections of z-stack confocal images of the TOMM20 fluorescence channel using ImageJ (Fiji) plug-in tool Mitochondria Analyzer [64]. The preprocessing parameters were set as follows: rolling (microns), 1,25 um; radius, 2; max slope, 2; and gamma, 0.8. The applied commands included subtracting the background, enhancing local contrast, and adjusting gamma. The threshold method used was the weighted mean method with a block size of 1.75 μm and a c value of 6. Postprocessing commands included removing outliers with a 2-pixel radius. At least four fields of view were analyzed for each biological replicate.

### 4.9 Flow cytometry

Cells were harvested using Accutase solution (#LM-T1735, Biosera), diluted in PBS with 3% fetal bovine serum (#FB-1101, Biosera) and 2 mM EDTA (#ED2SS, Sigma-Aldrich) and measured using CytoFLEX S flow cytometer (Beckman Coulter, Brea, CA, USA). Cell viability was measured using propidium iodide (#25535-16-4, P-lab, Prague, Czech Republic) in a final concentration of 600 nM. PI-positive (non-viable) cells were detected and quantified in the ECD (610/20) channel. To detect changes in mitochondrial membrane potential, cells were incubated with JC-1 dye (#65-0851-38, Invitrogen) for 10 minutes at 37 °C at a final concentration of 10 µg/ml. Cells treated with 10 uM FCCP (#HY100410, MedChemExpress, New Jersey, USA) for 1 h were used as a positive control. JC-1 fluorescence was excited by a 488 nm laser and detected using the FITC (525/40) and ECD (610/20) channels. To eliminate the spillover signal of JC-1 monomers to the ECD channel, fluorescence compensation was performed as previously described [64]. Gating strategies and representative analyses are provided in Supplementary Fig. 11.

### 4.10 Statistical analysis

Statistical analysis was performed in GraphPad Prism 8.0.2 software. For comparisons of multiple groups, one-way analysis of variance (ANOVA) with Dunnett’s test was used to compare the sample value with the corresponding control group. For comparing two groups, an unpaired t-test with Welch’s correction was used, assuming normal data distribution and similar variance between the compared groups. P values of p < 0.05 were considered statistically significant. *p < 0.05, **p < 0.01, ***p < 0.001, ****p < 0.0001. Results from at least 3 biological replicates were evaluated.

## AUTHOR CONTRIBUTIONS

A.K. performed most of the experiments and data analysis; M.S., M.M., and A.K. generated pCDH-CuO-MYC/MYCN vectors; A.K., T.B., and I.H. produced lentiviral particles; A.K. and I.H. established SHEP-CuO cell lines; I.H. performed immunoblotting analysis of MYC induction in pool populations and viability assays following tigecycline and chloramphenicol treatments. J. St. contributed to the generation of patient-derived neuroblastoma cell lines. J.Sk., A.K., and K.B. jointly conceived the main idea of this study. A.K. and J.Sk. designed the experiments, conceptualized and drafted the manuscript. All authors contributed to the discussion and revision of the manuscript. All authors read and approved the final version of the manuscript.

## CREDIT AUTHORSHIP CONTRIBUTION STATEMENT

**Adela Kubistova**: Conceptualization, Investigation, Methodology, Formal analysis, Data curation, Visualization, Writing – original draft. Writing – review & editing. **Iryna Horak**: Investigation, Methodology, Formal analysis, Writing – review & editing. **Tomas Barta**: Methodology, Resources, Writing – review & editing. **Marie Sulova**: Methodology, Writing – review & editing. **Martin Marek**: Methodology, Resources, Writing – review & editing. **Jaroslav Sterba:** Resources, Writing – review & editing. **Karolina Borankova**: Conceptualization, Writing – review & editing, **Jan Skoda**: Conceptualization, Supervision, Formal analysis, Funding acquisition, Project administration, Visualization, Writing – original draft. Writing – review & editing.

## ETHICS DECLARATION

The study was conducted according to the guidelines of the Declaration of Helsinki, and approved by the Masaryk University Research Ethics Committee (ref. no.: EKV-2024-031, granted on December 9, 2024) and the Ethics Board of the Faculty of Medicine, Masaryk University (ref. no.: 0299/2025, granted on March 14, 2025). All noncommercial patient-derived cell lines were obtained with written informed consent, including permission for secondary research use.

## FUNDING

This work was supported by the Czech Science Foundation (GA25-16629S) and the Ministry of Health of the Czech Republic in cooperation with the Czech Health Research Council under project no. NW26-07-00021.

## DECLARATION OF COMPETING INTEREST

The authors declare that they have no known competing financial interests or personal relationships that could have appeared to influence the work reported in this paper.

## ACKNOWLEDGEMENTS

A.K. acknowledges support from the Grant Agency of Masaryk University (MUNI/C/1790/2023). The work of M.M. on cumate-inducible vectors was supported by the Czech Science Foundation (GX25-17329X). J.Sk. was supported by the Ministry of Health of the Czech Republic in cooperation with the Czech Health Research Council within the National Institute for Cancer Research, project No. NW26A-NUVR. We are thankful to Dr. Tomáš Loja for his technical assistance with fluorescence-assisted cell sorting and acknowledge the Core Facility Genomics (CEITEC, Brno, Czech Republic) for access to their services.

## SUPPLEMENTARY MATERIAL

### SUPPLEMENTARY FIGURES

**Supplementary Figure 1.**
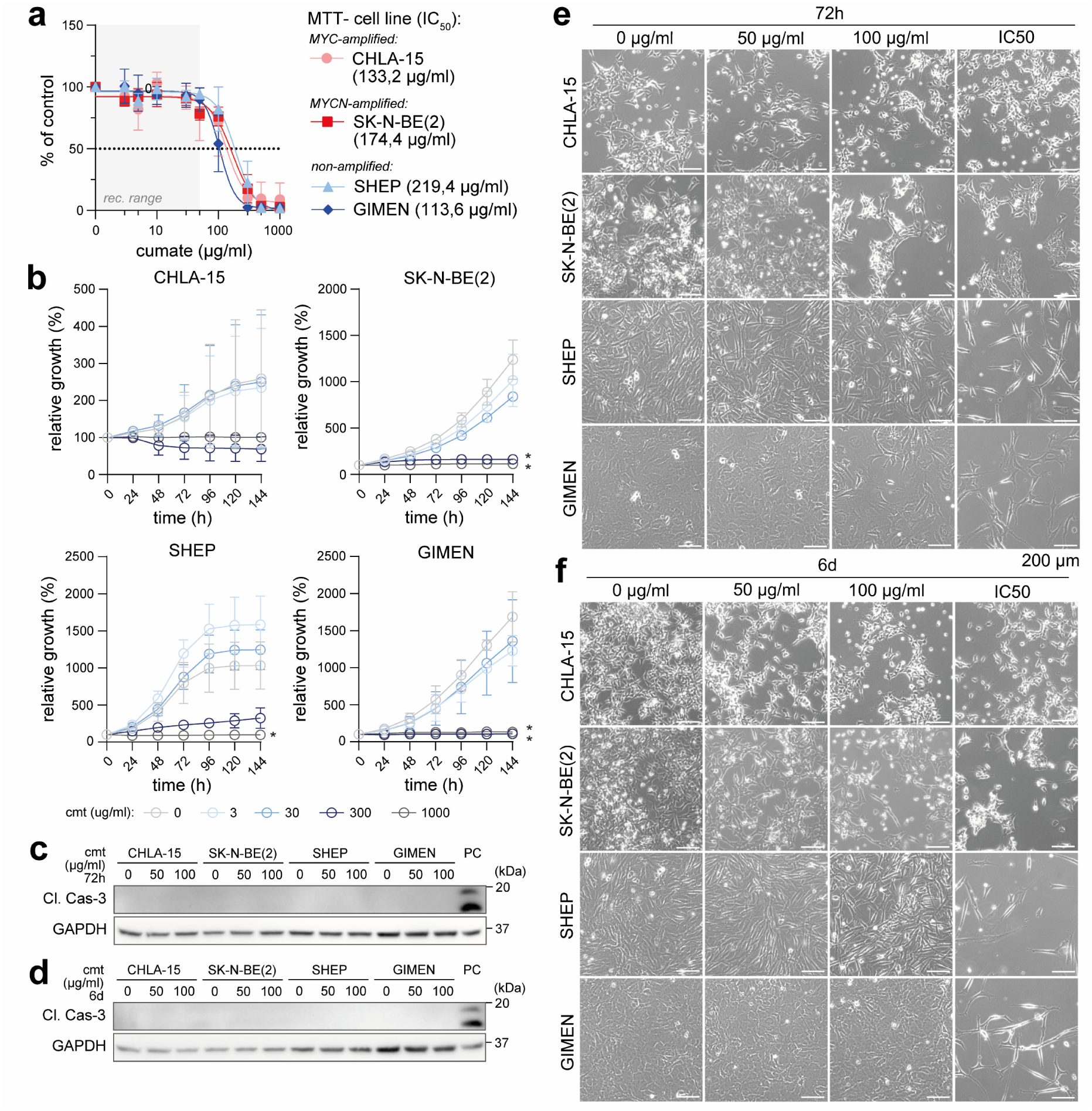
The maximum recommended dose of cumate remains non-perturbing during prolonged treatment. (**a**) Sensitivity of neuroblastoma cell lines to cumate tested by MTT assay after 6 d of treatment (calculated IC50 values are indicated). Data are presented as mean ± SD (biological n = 3, technical n = 3). (**b**) Live-cell imaging growth rate analysis of neuroblastoma cell lines after 6 d of cumate treatment. Data are presented as mean ± SD (biological n = 3, technical n = 3). Representative images of neuroblastoma cells after 72 h. (**c,d**) Western blot detection of activated caspase-3 protein levels in a panel of neuroblastoma cells after 72 h (**c**) and 6 d (**d**) of cumate treatment at the indicated concentrations (biological n = 3). Densitometric analysis was not performed due to the absence of a signal. PC, positive control (CHLA-15 cells, 24 h treatment with 50 µM doxycycline). (**e,f**) Cell morphology after 3 d (**e**) and 6 d (**f**) of cumate treatment, demonstrating no detectable morphological changes at concentrations up to 100 µg/ml. Statistical significance was determined by one-way ANOVA followed by Dunnett’s multiple comparisons test (**b**), *p < 0.05.

**Supplementary Figure 2.**
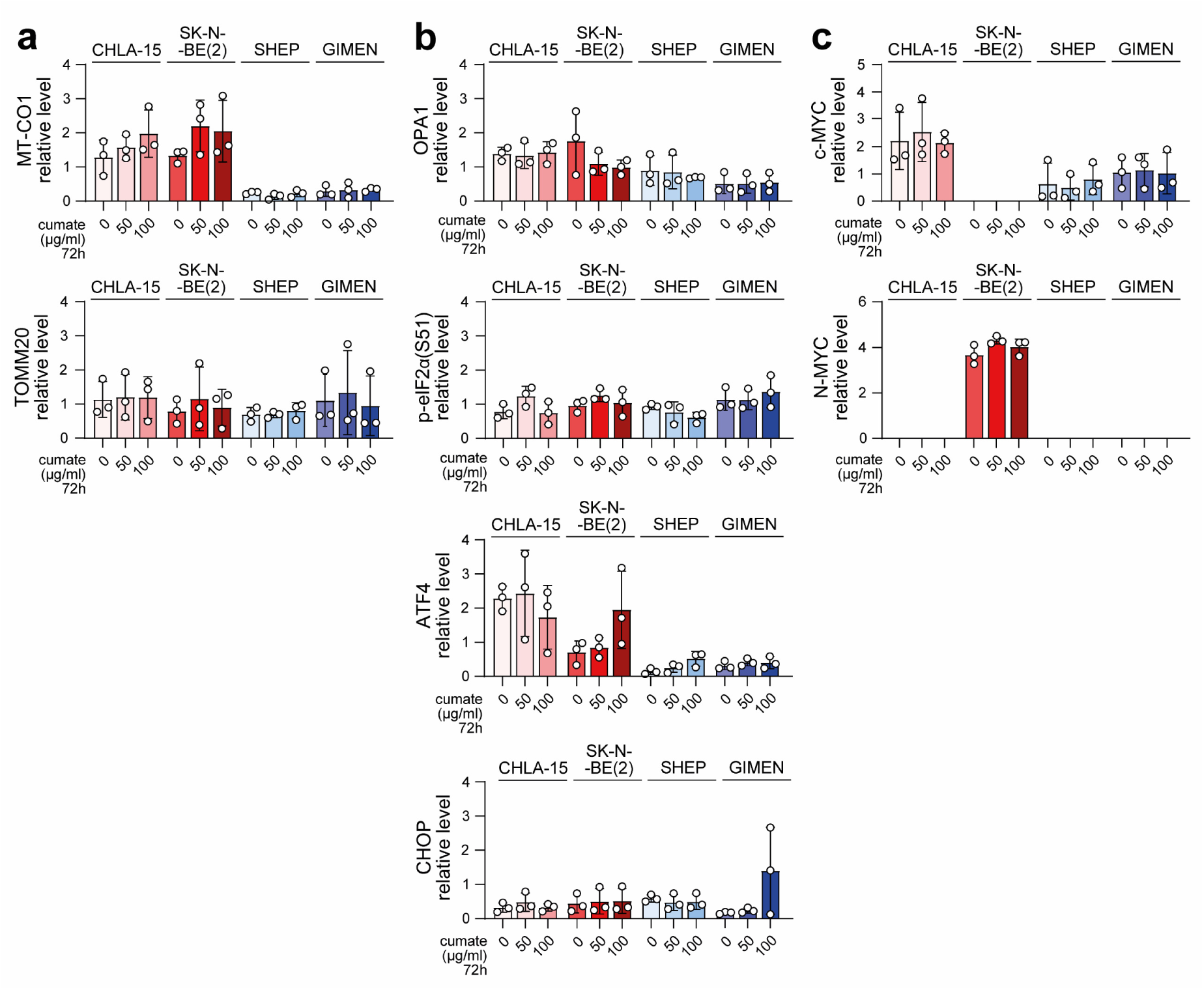
Cumate does not affect levels of mitochondrial proteins, mitochondrial stress markers, or MYC proteins. Densitometric analysis of western blotting detection of mitochondrial proteins (**a**), markers of ISR (**b**), and MYC proteins (**c**) after 72 h of cumate treatment at the indicated concentrations. Normalized protein levels are plotted relative to the average of all samples (mean ± SD, biological n = 4). Statistical significance was determined by one-way ANOVA followed by Dunnett’s multiple comparisons test, *p < 0.05.

**Supplementary Figure 3.**
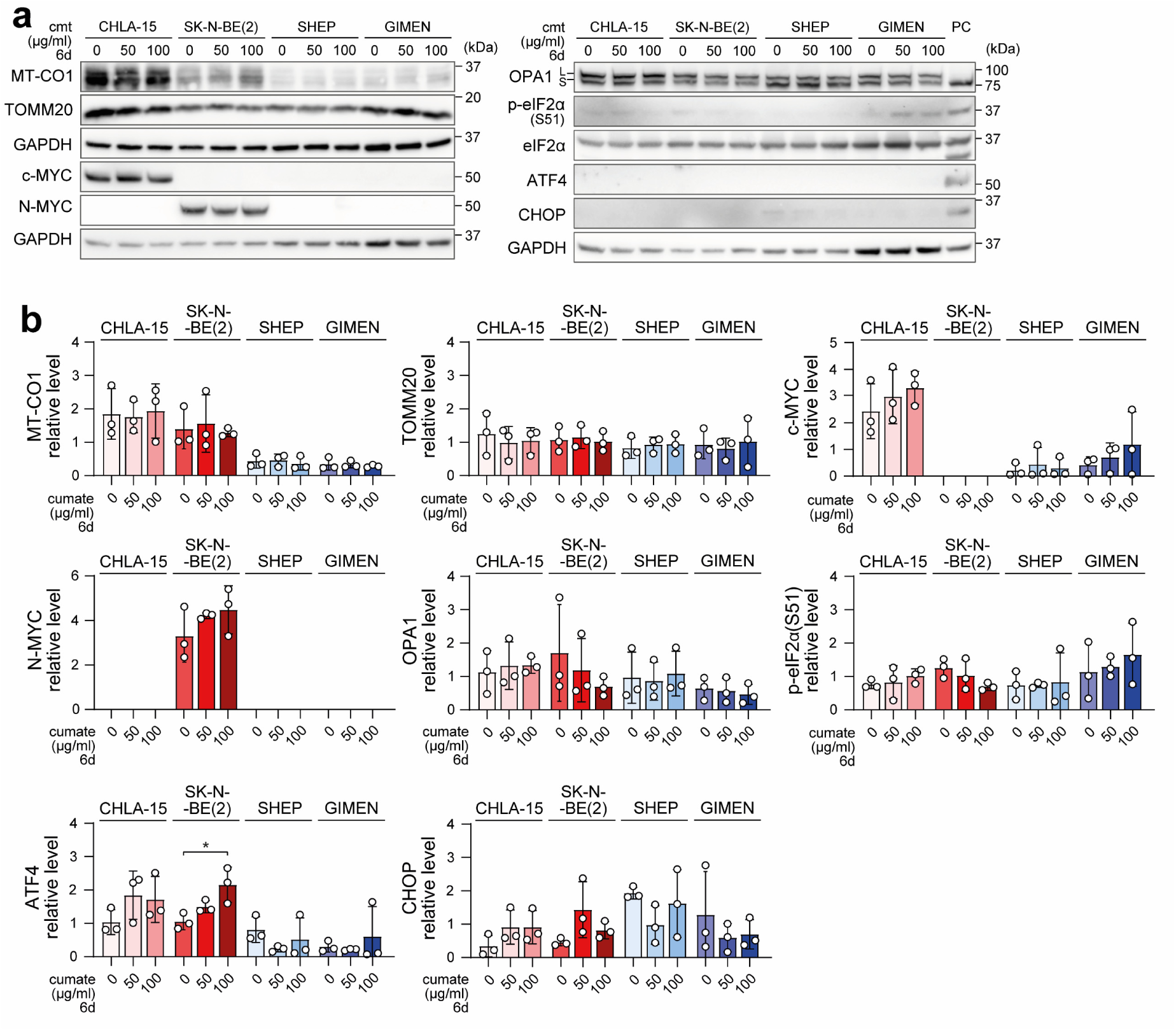
Prolonged cumate treatment does not alter mitochondrial or MYC protein levels. Western blot detection (**a**) and densitometric analysis (**b**) of mitochondrial proteins, markers of ISR and MYC proteins after 6 d of cumate treatment at indicated concentrations. Normalized protein levels are plotted relative to the average of all samples (mean ± SD, biological n = 4). Statistical significance was determined by one-way ANOVA followed by Dunnett’s multiple comparisons test (**b**), *p < 0.05.

**Supplementary Figure 4.**
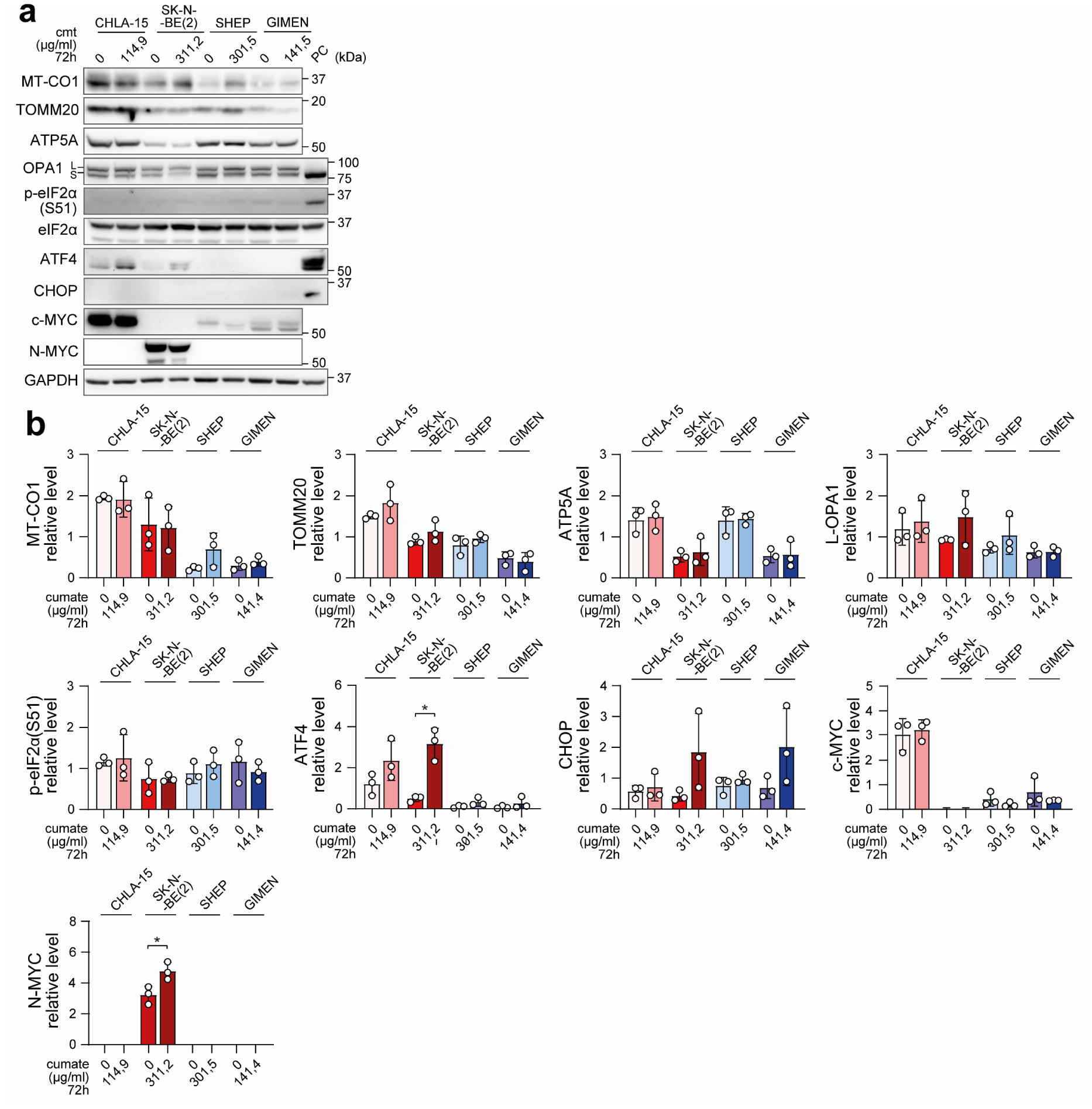
Expression of key proteins following short-term exposure to high-dose cumate. Western blot detection (**a**) and densitometric analysis (**b**) of mitochondrial proteins, markers of ISR and MYC proteins after 72 h of cumate treatment at IC_50_ concentrations calculated for each cell line by MTT assay (Fig. 1b, S1a). Normalized protein levels are plotted relative to the average of all samples (mean ± SD, biological n = 4). PC, positive control (CHLA-15 cells, 24 h treatment with 50 µM doxycycline). Statistical significance was determined by unpaired two-tailed t-test with Welch correction (**b**), *p < 0.05.

**Supplementary Figure 5.**
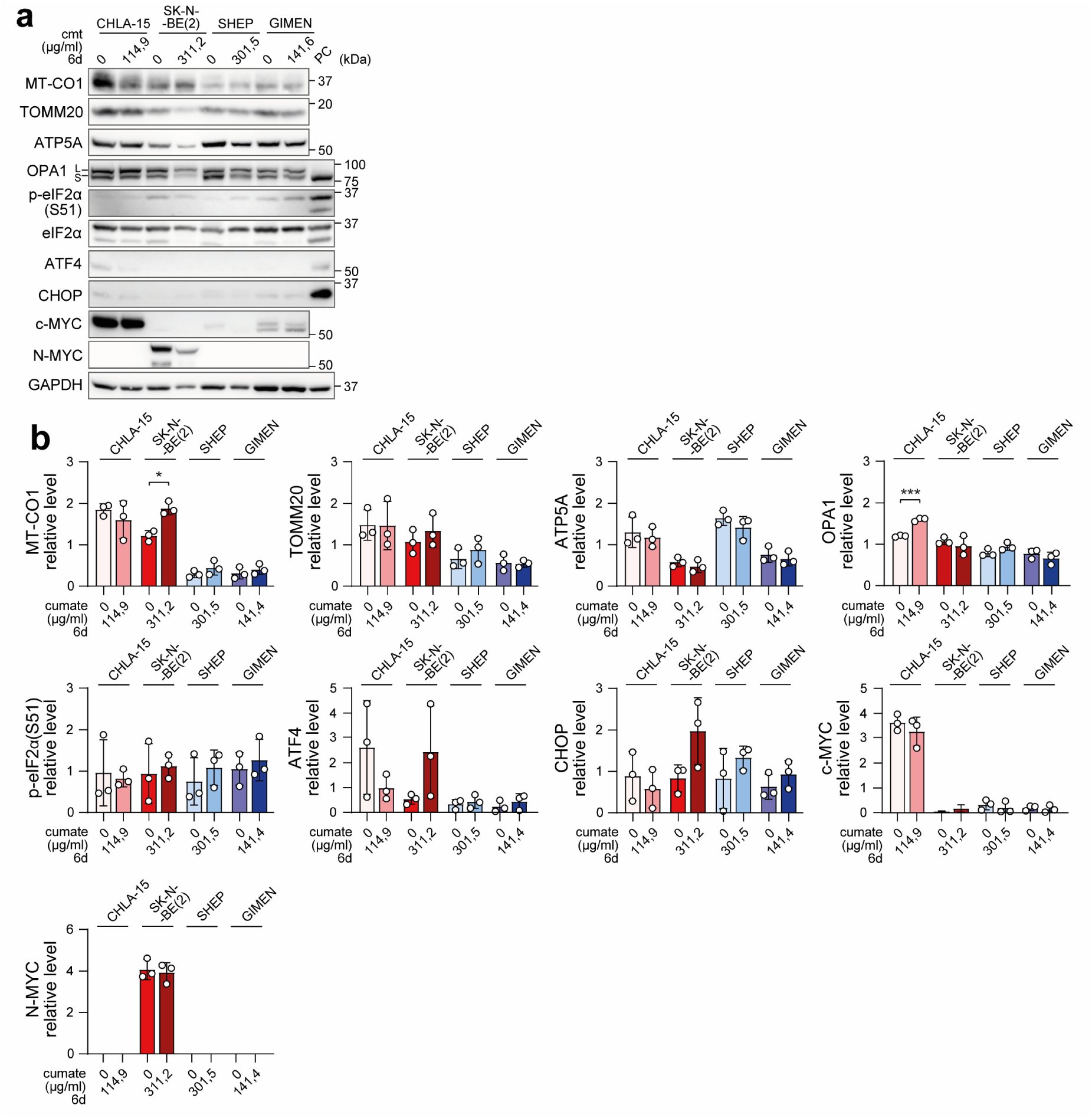
Expression of key proteins after prolonged exposure to high-dose cumate. Western blot detection (**a**) and densitometric analysis (**b**) of mitochondrial proteins, markers of ISR and MYC proteins after 6 d of cumate treatment at IC_50_ concentrations calculated for each cell line by MTT assay (Fig. 1b, S1a). Normalized protein levels are plotted relative to the average of all samples (mean ± SD, biological n = 4). PC, positive control (CHLA-15 cells, 24 h treatment with 50 µM doxycycline). Statistical significance was determined by unpaired two-tailed t-test with Welch correction (**b**), *p < 0.05, **p < 0.01, ***p < 0.001.

**Supplementary Figure 6.**
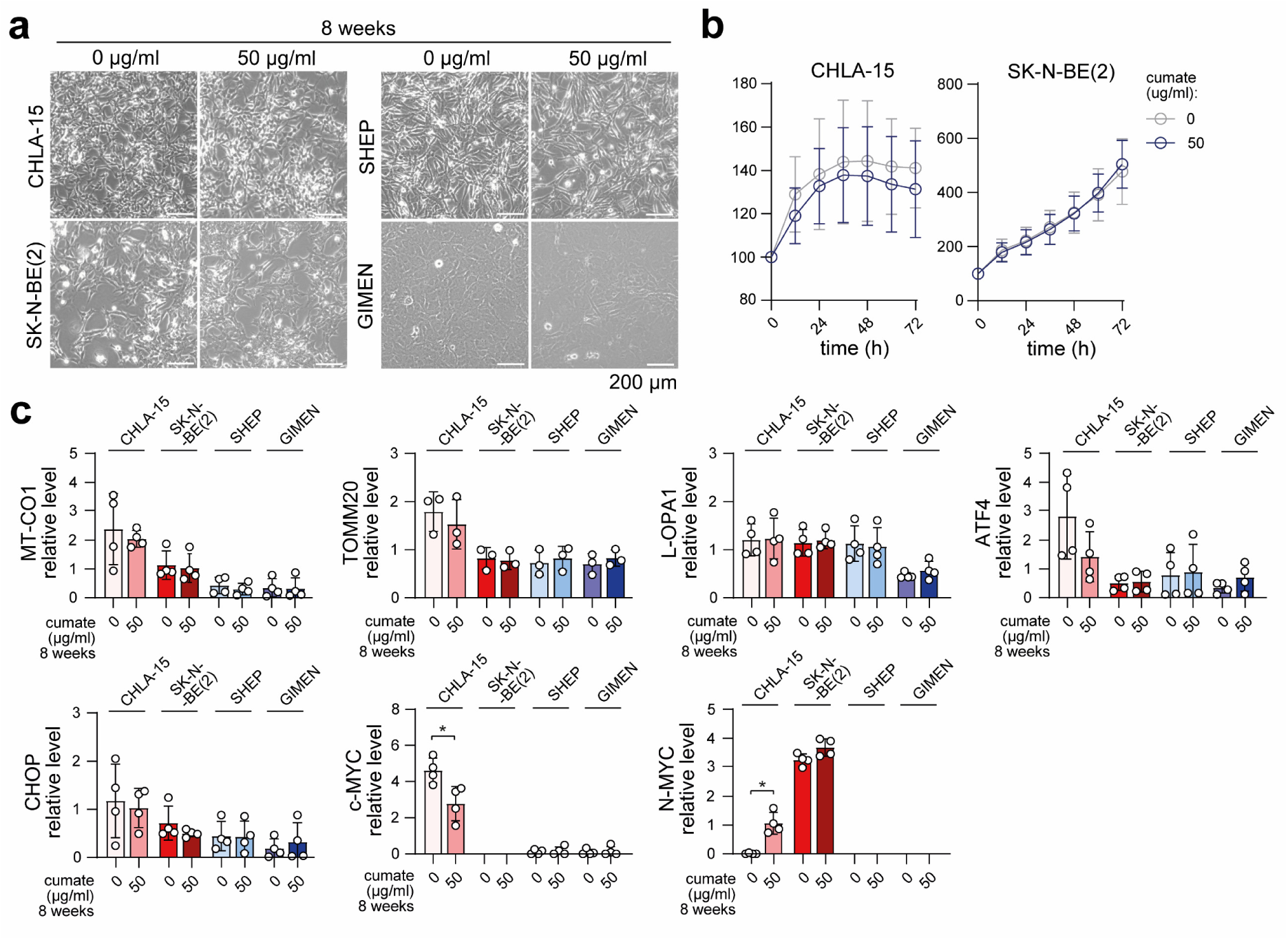
Long-term treatment with low doses of cumate does not significantly affect neuroblastoma cells. (**a**) Representative images of neuroblastoma cells after 8 weeks of 50 µg/ml cumate treatment, demonstrating no detectable morphological or proliferatory changes. (b) Live-cell imaging growth rate analysis of neuroblastoma cell lines after 8 weeks of 50 µg/ml cumate treatment. Data are presented as mean ± SD (biological n = 3, technical n = 3). (**c**) Densitometric analysis of mitochondrial proteins, markers of ISR and MYC proteins after 8 weeks of cumate treatment. Statistical significance was determined by unpaired two-tailed t-test with Welch correction (**b,c**), *p < 0.05.

**Supplementary Figure 7.**
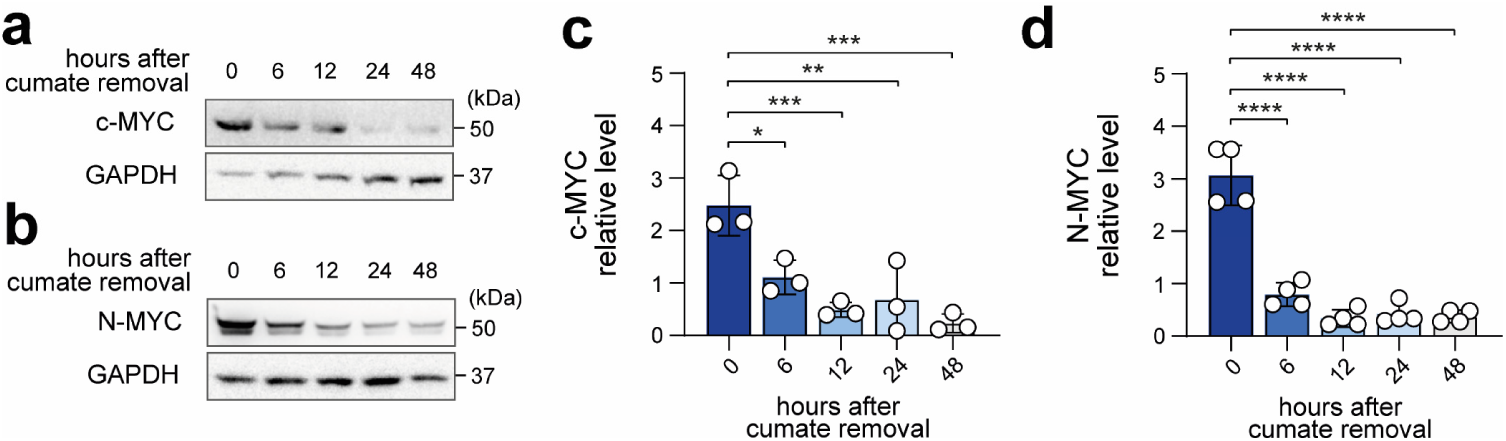
Cumate-induced c-MYC and N-MYC expression is reversible upon removal of cumate from the cell culture medium. Western blot detection (**a,b**) and densitometric analysis (**c,d**) of c-MYC and N-MYC protein levels following treatment with 10 µg/ml cumate for 24 h, followed by replacement of cumate-containing medium with fresh cumate-free medium. Normalized protein levels are plotted relative to the average of all samples (mean ± SD, biological n = 3–4). Statistical significance was determined by one-way ANOVA followed by Dunnett’s multiple comparisons test (**a,b**), *p < 0.05, **p < 0.01, ***p < 0.001, ***p < 0.0001.

**Supplementary Figure 8.**
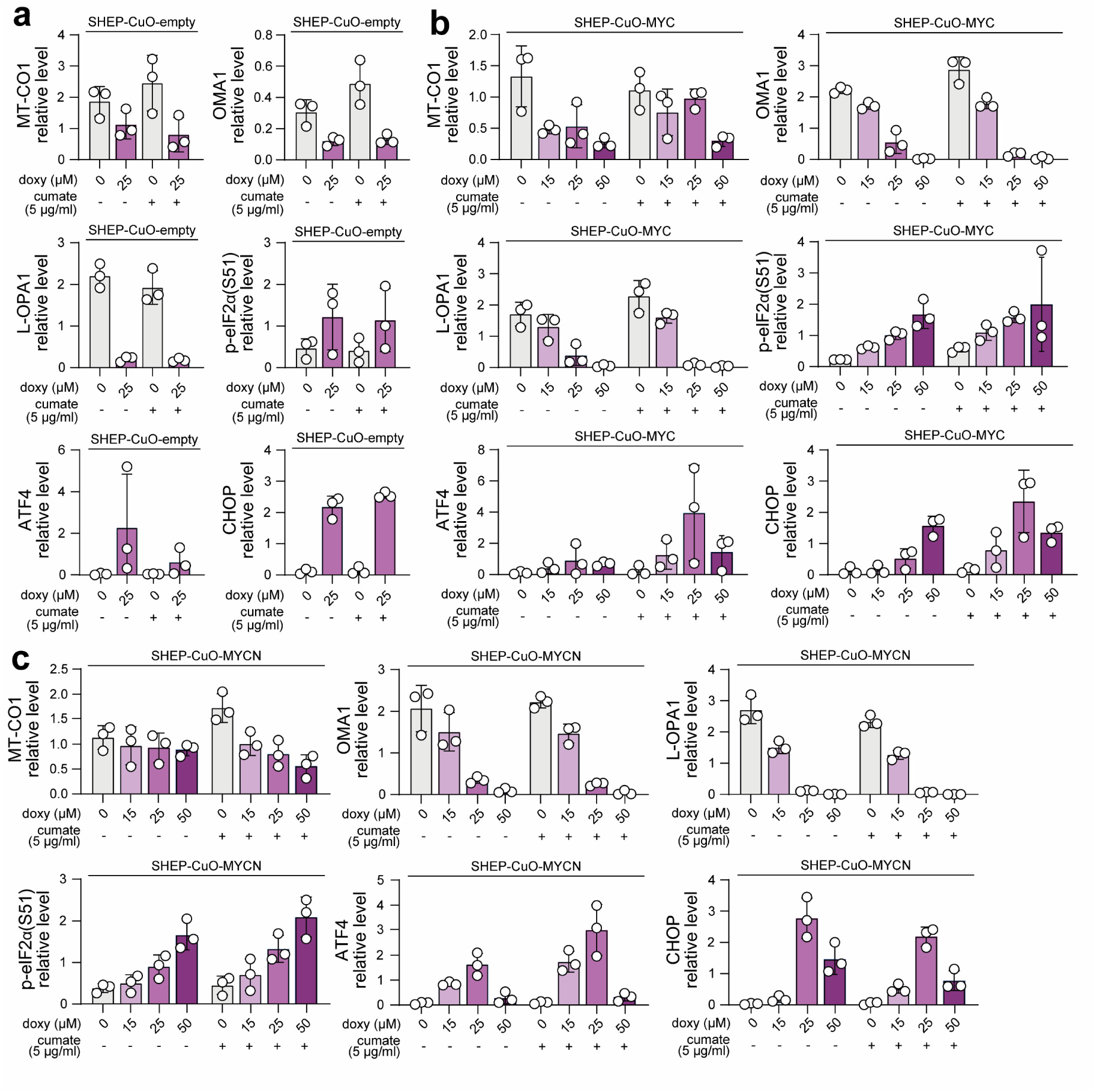
Doxycycline induces comparable activation of mitoISR in MYC-off and MYC-on conditions. Densitometric analysis of western blotting detection of mitochondrial proteins, markers of ISR and MYC proteins in single-cell clone-derived SHEP-CuO-empty (**a**), SHEP-CuO-MYC (**b**) and SHEP-CuO-MYCN (**c**) cell lines after at least 24 h of treatment with 5 ug/ml cumate followed by 24 h of treatment with indicated doxycycline concentrations. Normalized protein levels are plotted relative to the average of all samples (mean ± SD, biological n = 3, each point represents an individual single-cell clone). Statistical significance was determined by unpaired two-tailed t-test with Welch correction. The same doxycycline concentrations in the presence and absence of cumate were compared to determine the specific effect of MYC-on conditions, *p < 0.05.

**Supplementary Figure 9.**
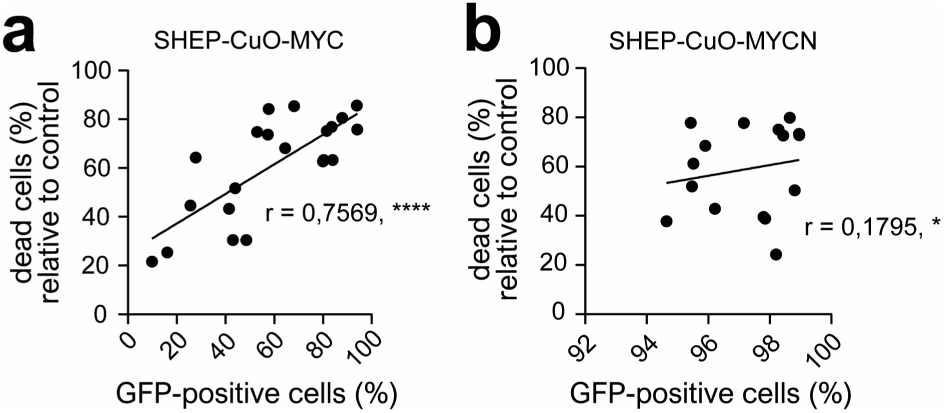
Doxycycline-induced cell death correlates with the induction of MYC levels. Correlation analysis between the percentage of dead cells in SHEP-CuO-MYC (**a**) and SHEP-CuO-MYCN (**b**) after 24 h treatment with 75 μM doxycycline and the percentage of GFP-positive cells in untreated controls, r = Pearson correlation coefficient (biological n ≥ 16; single-cell clones and pool populations).

**Supplementary Figure 10.**
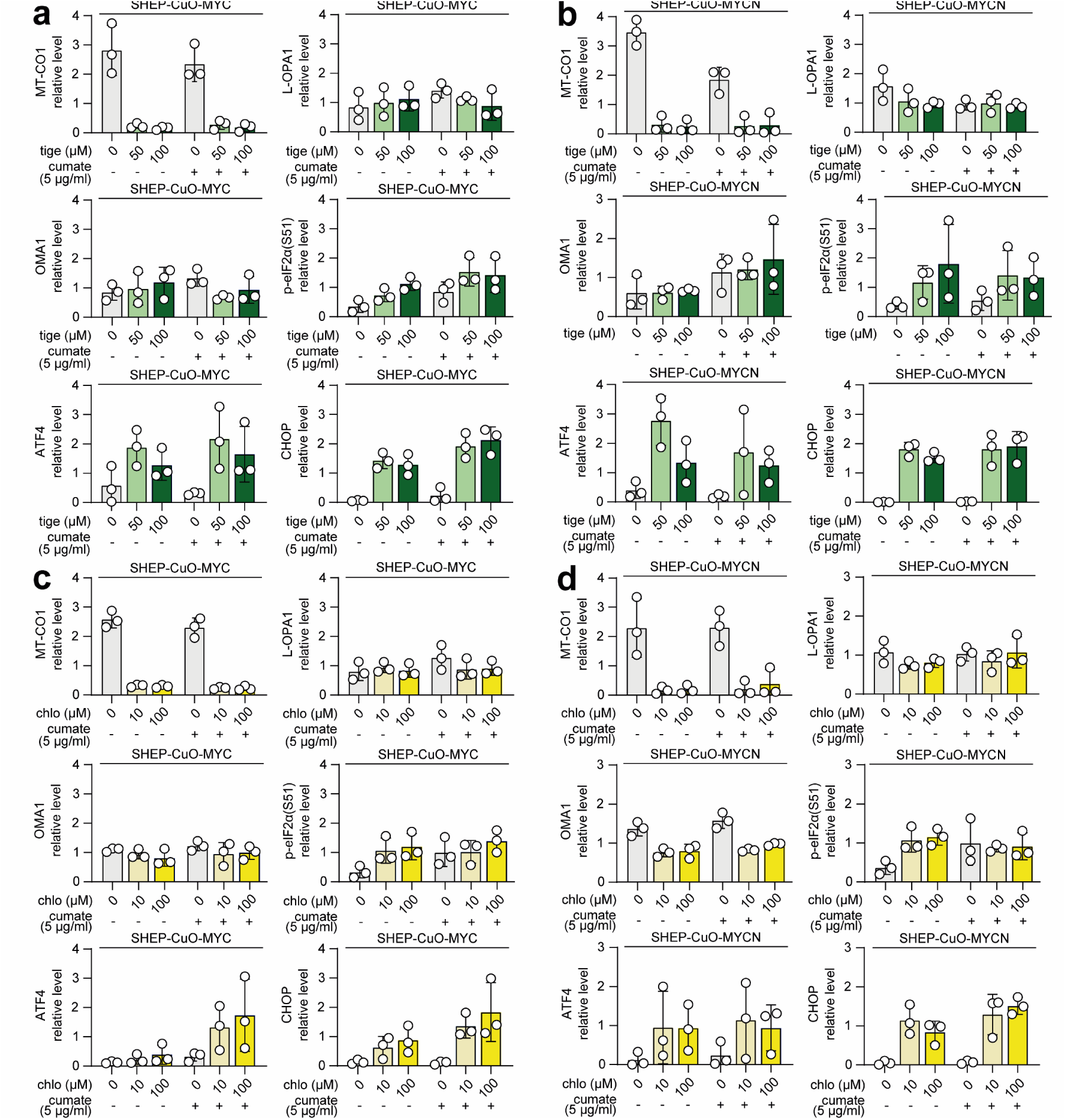
MitoISR activation by ribosomal antibiotics. Densitometric analysis of western blotting detection of mitochondrial proteins, markers of ISR and MYC proteins in selected single-cell clone-derived SHEP-CuO-MYC (**a,c**) and SHEP-CuO-MYCN (**b,d**) cell lines after at least 24 h of treatment with 5 µg/ml cumate followed by 24 h of treatment with indicated tigecycline (**a,b**) or chloramphenicol (**c,d**) concentrations. Normalized protein levels are plotted relative to the average of all samples (mean ± SD, biological n = 3, each point represents an individual single-cell clone). Statistical significance was determined by unpaired two-tailed t-test with Welch correction. The same drug concentrations in the presence and absence of cumate were compared to determine the specific effect of MYC-high conditions, *p < 0.05.

**Supplementary Figure 11.**
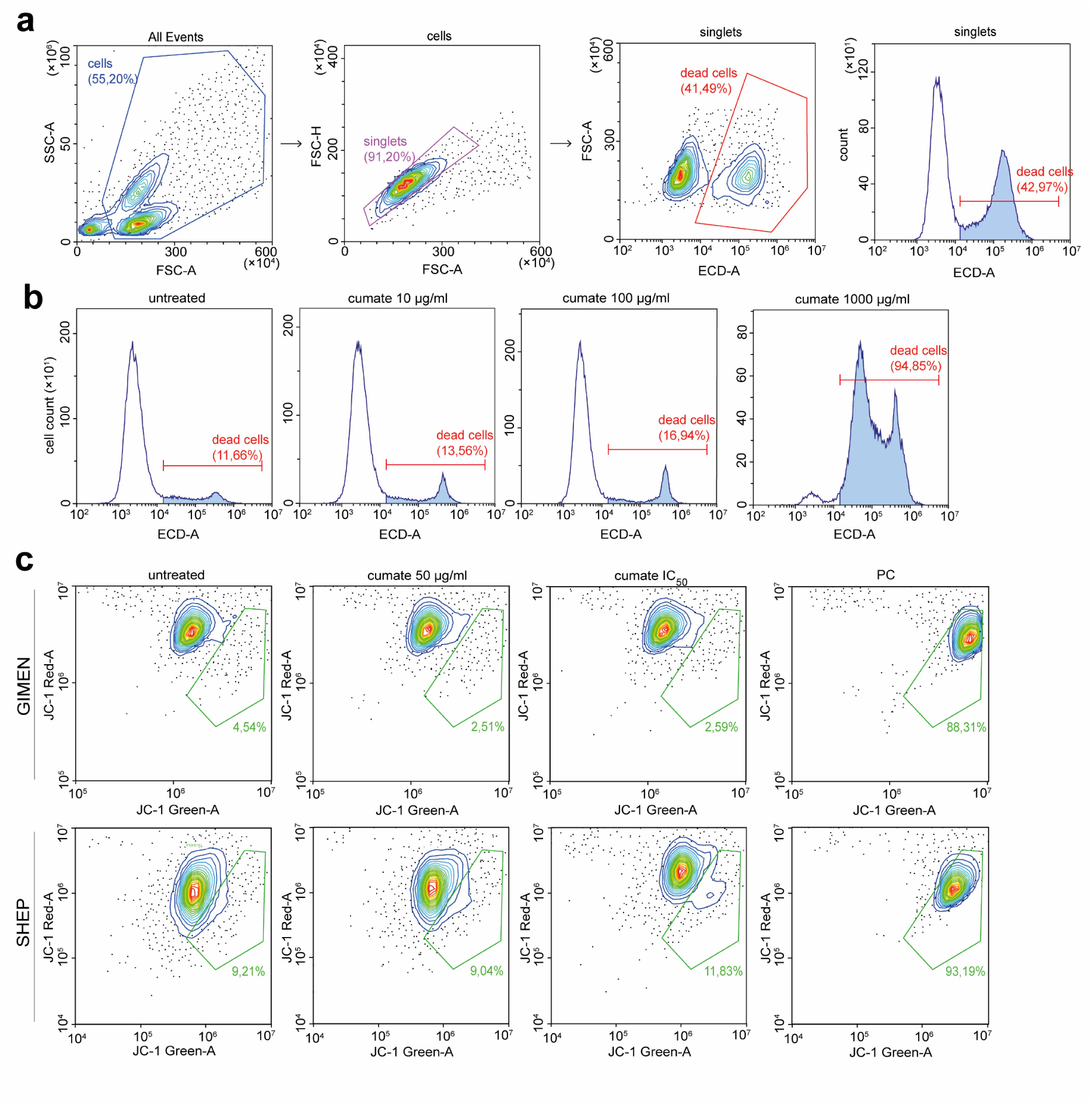
Flow cytometry gating strategy and representative analyses of dead cells and mitochondrial depolarization. (**a**) Flow cytometry gating strategy for detection of dead cells. Debris was excluded based on FSC-A vs SSC-A parameters. Singlets were selected using FSC-A vs FSC-H gating to remove doublets. Dead cells were identified by PI staining using ECD-A vs FSC-A, with PI-positive events classified as dead cells. (**b**) Representative analysis of cell death following cumate treatment at the indicated concentrations. Gating thresholds were defined using positive control, cells heated to 70 °C for 15 minutes. (**c**) Representative analysis of changes in mitochondrial depolarization following cumate treatment at the indicated concentrations using JC-1 staining. Gating thresholds were defined using positive control (PC), 1h treatment with 10 µM FCCP.

### SUPPLEMENTARY TABLES

**Supplementary Table 1.**
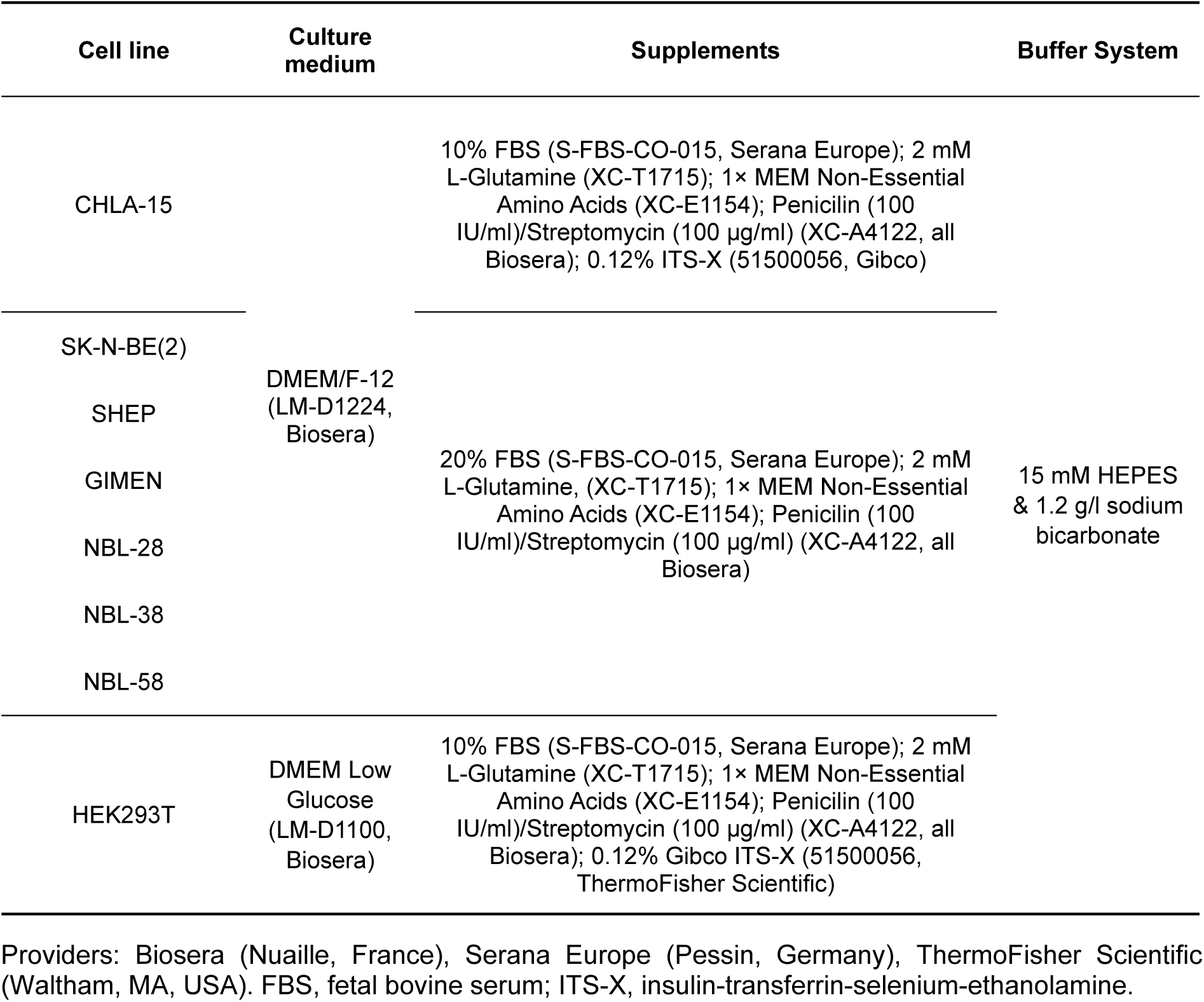
Summary of cell culture media composition.

**Supplementary Table 2.**
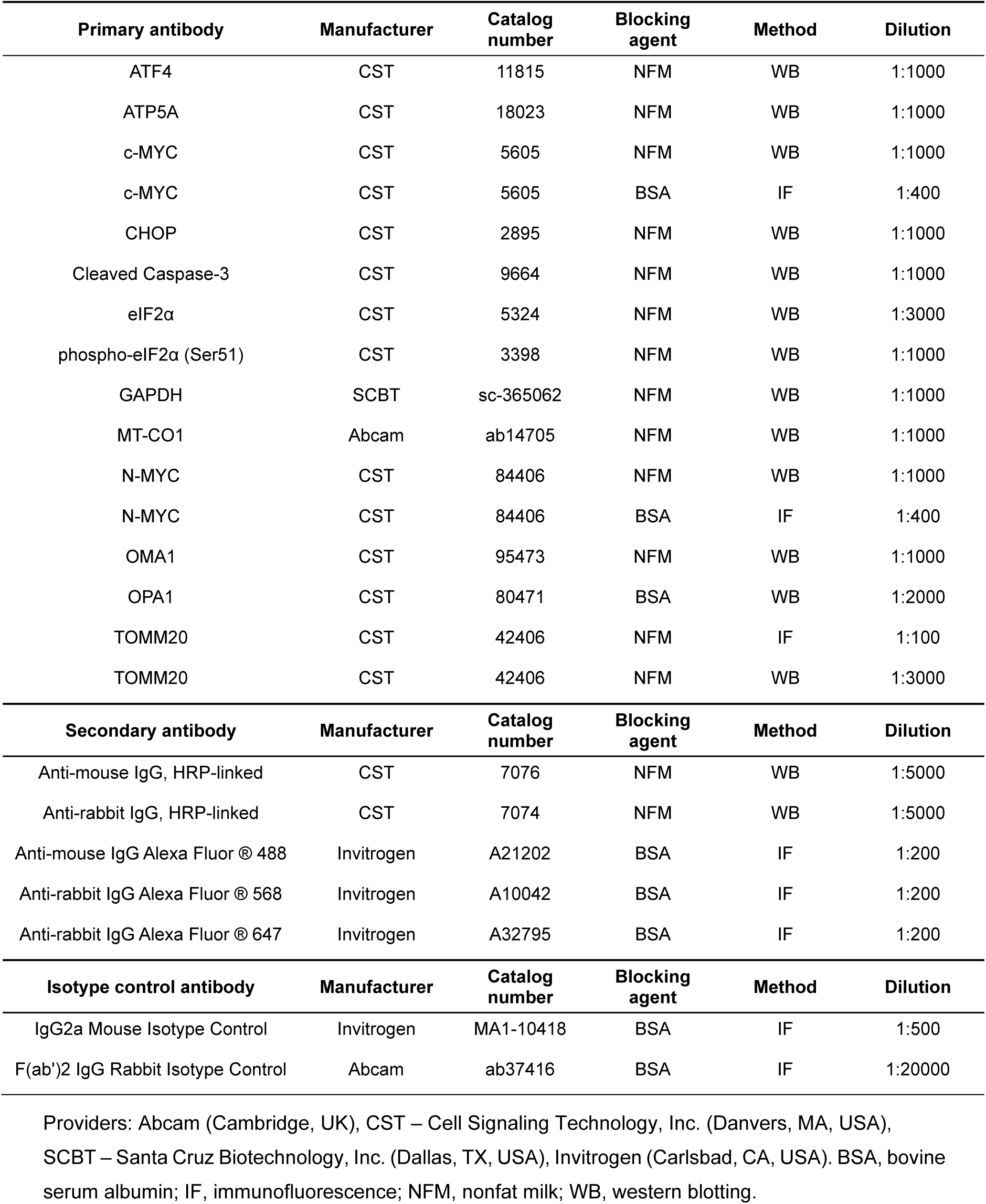
Antibodies used for immunodetection.

Providers: Biosera (Nuaille, France), Serana Europe (Pessin, Germany), ThermoFisher Scientific (Waltham, MA, USA). FBS, fetal bovine serum; ITS-X, insulin-transferrin-selenium-ethanolamine.
| Primary antibody | Manufacturer | Catalog number | Blocking agent | Method | Dilution |
| --- | --- | --- | --- | --- | --- |
| ATF4 | CST | 11815 | NFM | WB | 1:1000 |
| ATP5A | CST | 18023 | NFM | WB | 1:1000 |
| c-MYC | CST | 5605 | NFM | WB | 1:1000 |
| c-MYC | CST | 5605 | BSA | IF | 1:400 |
| CHOP | CST | 2895 | NFM | WB | 1:1000 |
| Cleaved Caspase-3 | CST | 9664 | NFM | WB | 1:1000 |
| eIF2α | CST | 5324 | NFM | WB | 1:3000 |
| phospho-eIF2α (Ser51) | CST | 3398 | NFM | WB | 1:1000 |
| GAPDH | SCBT | sc-365062 | NFM | WB | 1:1000 |
| MT-CO1 | Abcam | ab14705 | NFM | WB | 1:1000 |
| N-MYC | CST | 84406 | NFM | WB | 1:1000 |
| N-MYC | CST | 84406 | BSA | IF | 1:400 |
| OMA1 | CST | 95473 | NFM | WB | 1:1000 |
| OPA1 | CST | 80471 | BSA | WB | 1:2000 |
| TOMM20 | CST | 42406 | NFM | IF | 1:100 |
| TOMM20 | CST | 42406 | NFM | WB | 1:3000 |
| Secondary antibody | Manufacturer | Catalog number | Blocking agent | Method | Dilution |
| Anti-mouse IgG, HRP-linked | CST | 7076 | NFM | WB | 1:5000 |
| Anti-rabbit IgG, HRP-linked | CST | 7074 | NFM | WB | 1:5000 |
| Anti-mouse IgG Alexa Fluor $\text{\textcircled{R}}$ 488 | Invitrogen | A21202 | BSA | IF | 1:200 |
| Anti-rabbit IgG Alexa Fluor $\text{\textcircled{R}}$ 568 | Invitrogen | A10042 | BSA | IF | 1:200 |
| Anti-rabbit IgG Alexa Fluor $\text{\textcircled{R}}$ 647 | Invitrogen | A32795 | BSA | IF | 1:200 |
| Isotype control antibody | Manufacturer | Catalog number | Blocking agent | Method | Dilution |
| IgG2a Mouse Isotype Control | Invitrogen | MA1-10418 | BSA | IF | 1:500 |
| F(ab')<sub>2</sub> IgG Rabbit Isotype Control | Abcam | ab37416 | BSA | IF | 1:20000 |

Providers: Abcam (Cambridge, UK), CST – Cell Signaling Technology, Inc. (Danvers, MA, USA), SCBT – Santa Cruz Biotechnology, Inc. (Dallas, TX, USA), Invitrogen (Carlsbad, CA, USA). BSA, bovine serum albumin; IF, immunofluorescence; NFM, nonfat milk; WB, western blotting.

**Supplementary Table 3.**
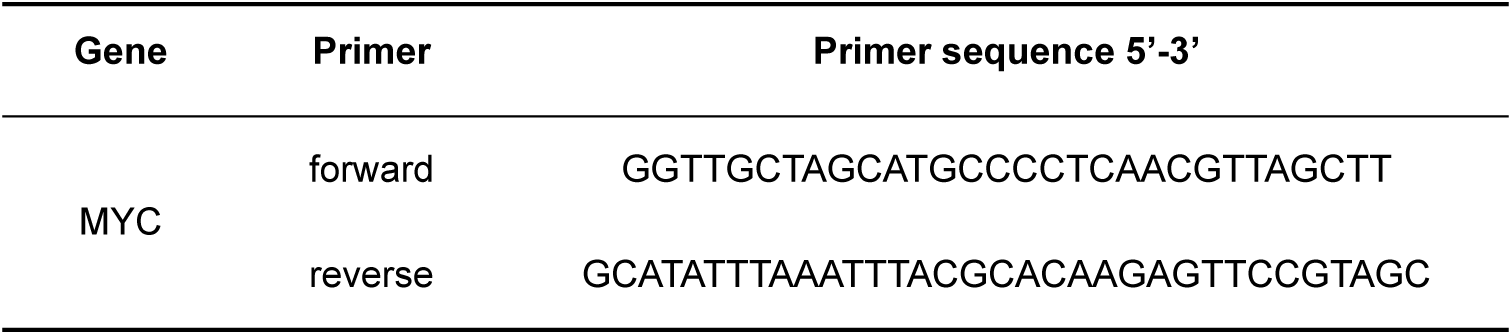
Cloning primer sequences.

## Notes

### Competing Interest Statement

The authors have declared no competing interest.

### Summary of Updates

Results section expanded to include preclinical validation in patient-derived cell lines, parts of the manuscript rewritten accordingly; Added Figure 6; Authors list expanded.

